# Antibiotic potentiators from microbiota that permeate the *Pseudomonas aeruginosa* cell envelope

**DOI:** 10.64898/2026.11.23.768654

**Authors:** Rahgavi Poopalarajah, Andrew J. Flewelling, Shirin Afroj, Noah B. Toppings, Ben Rowley, Kevin C. Sule, Nathan K. McCartney, Andrew J. Geppert, Vanessa Adams, Yanqi Li, Carina Thomas, Shannon Pyke, Emma J. Griffiths, Nithu Sara John, Charlotte Barclay, Rhiannon Cameron, Damion Dooley, Barbara Waddell, Craig Brideau, Zajeba Tabashsum, Jamie D. Liu, Michelle Angeles-Solano, Michael D. Parkins, John M. Conly, Ayush Kumar, Pina Colarusso, Elmar J. Prenner, William Hsiao, Kathy D. McCoy, Douglas J. Mahoney, Sarah E. Rowe, Darren J. Derksen, Joe J. Harrison

**Affiliations:** Department of Biological Sciences, University of Calgary, 2500 University Drive NW, Calgary, AB, Canada, T2N 1N4; One Health, University of Calgary, 3280 Hospital Drive NW, Calgary, AB, Canada T2N 4Z6; Department of Chemistry, University of Calgary, 2500 University Dr NW, Calgary, AB, Canada, T2N 1N4; Department of Microbiology, University of Manitoba, Winnipeg, MB, Canada, R3T 2N2; Department of Physiology and Pharmacology, University of Calgary, 3330 Hospital Drive NW, Calgary, AB, Canada, T2N 4N1; Centre for Infectious Disease Genomics and One Health, Faculty of Health Sciences, Simon Fraser University, Burnaby, BC, Canada, V5A 1S6; Department of Medicine, University of Calgary, 3330 Hospital Drive NW, Calgary, AB, Canada T2N 4N1; Department of Microbiology, Immunology and Infectious Diseases, University of Calgary, 3330 Hospital Drive NW, Calgary, AB, Canada T2N 4N1; Snyder Institute for Chronic Diseases, 3280 Hospital Drive NW, Calgary, AB, Canada T2N 4Z6; Department of Microbiology and Immunology, University of North Carolina, Chapel Hill, North Carolina, USA 27599

**Keywords:** antimicrobial resistance, antibiotics, carbapenem, antibiotic potentiator, natural products, microbiome, bacteria, *Pseudomonas aeruginosa*, membrane permeability, fatty amide

## Abstract

Antibiotic potentiators are bioactive substances that enhance antibiotic activity. Many antibiotics have originated from natural product screens; however, systematic microbial testing that has powered antibiotic discovery has seldom been applied to potentiators. Here we describe a natural product platform based on medium-throughput parallel microbial culture to identify antibiotic potentiators. Using carbapenem-resistant *Pseudomonas aeruginosa* exhibiting resistance due to reduced drug uptake, we identified many bacterial species producing spent media that potentiate last-resort antibiotics. Chemical analyses identified that one group of bioactive molecules was *cis*-unsaturated fatty amides. Biochemical, genetic, and structure-activity relationship studies showed that these substances increase small molecule uptake by altering fluidity and compromising the permeability barrier of bacterial membranes. *Cis*-unsaturated fatty amides with imipenem reduced the load of carbapenem-resistant *P. aeruginosa* in a murine wound infection model. We propose that our methods can drive the discovery of natural antibiotic potentiators and advance an understanding of biotic factors influencing antimicrobial resistance.

## INTRODUCTION

Antimicrobial resistance is a global health and development threat^1,2^. Antimicrobials are broadly defined as substances that can kill microorganisms (such as protozoa, fungi, bacteria and viruses) or inhibit their growth. Antibiotics are natural, semi-synthetic and synthetic antimicrobials that are active against bacteria and used to treat or prevent infections in humans and/or animals. In 2021, 4.71 million people globally lost their lives to 22 different bacterial pathogens resistant to at least 1 antibiotic^3^, and at least 1.14 million of these deaths were directly attributable to antibiotic resistance^4^. So, antibiotic resistance killed ∼1 person every 30 seconds.

The Gram-negative opportunistic pathogen *Pseudomonas aeruginosa* – one of the leading causes of life-endangering bacterial infections worldwide^5^ – has evolved many means to resist antibiotics^6^. At a molecular level, one mechanism involves efflux pumps, which expel antibiotics from cells^7–9^. Other mechanisms include enzymes that modify or degrade antibiotics, changes to the targets inside cells that nullify antibiotic mechanism-of-action, and adaptations that affect the permeability of cell membranes to antibiotics^7–9^.

One strategy to manage antibiotic resistant infections has been to develop potentiators^9–11^. These substances may have little or no antimicrobial toxicity alone but act to interfere with resistance mechanisms or augment antibiotic activity^9,11^. Some antibiotic potentiators target host defense mechanisms that can synergize with antibiotics^9^; others directly counter antibiotic resistance by blocking efflux pumps, inhibiting antibiotic-modifying enzymes, or permeabilizing membranes^9^.

The cell envelope of diderms like *P. aeruginosa*, which is comprised of the peptidoglycan layer and inner and outer membranes, provides considerable protection from toxic compounds. The outer membrane – composed of an asymmetric bilayer of tightly packed lipopolysaccharide (LPS) and phospholipid in outer and inner leaflets, respectively – is a formidable permeability barrier^12–14^. The outer membrane excludes many large and hydrophobic molecules^12^, constraining the influx of some antibiotics, like carbapenems, to porins^15^. By contrast, the inner membrane limits hydrophilic molecules from crossing into the cytoplasm^16,17^.

The World Health Organization identifies carbapenem-resistant *P. aeruginosa* as a public health threat^5,18^. While some of these bacteria acquire genes for carbapenem-degrading metallo-β-lactamases, many others acquire resistance via mutations in the porins OprD (also called OccD1 or PA0958) and OpdP (OccD3 or PA4501)^19,20^. These mutations are particularly common among carbapenem-resistant *P. aeruginosa* isolates in the Asia-Pacific region^21^, and in people with Cystic Fibrosis that are afflicted with chronic airways infections^22^. Conventional small molecule libraries mostly contain compounds with physical and chemical properties that are incompatible with outer membrane permeability^15^. For this reason, innovative screening techniques are necessary to drive discovery of antibiotic potentiators that can breach the *P. aeruginosa* cell envelope.

While most antibiotics are derived from microbial natural products^23–25^, only a few antibiotic potentiators that inhibit antibiotic-degrading enzymes have originated from these sources^26–29^. Nevertheless, the systematic testing of microbes that was used to drive antibiotic discovery in the 1940s^24^ has rarely been applied to potentiator discovery. This knowledge gap likely remains poorly explored due to the technical challenge of identifying antibiotic-sensitive, potentiator-producing organisms in antibiotic-amended co-cultures with antibiotic-resistant organisms.

In this work, we describe a culture-dependent screening method to identify microbes producing antibiotic potentiators in their spent media. This method provides access to natural chemical space unrepresented in synthetic, small molecule libraries. We identified dozens of bacterial species producing meropenem potentiators by probing a *P. aeruginosa* strain with loss-of-function mutations in the gene for the OprD uptake porin. Several species produced a fatty amide, oleamide, that potentiated multiple classes of antibiotics against *P. aeruginosa*. Other *cis*-unsaturated fatty amides (palmitoleamide, vaccenamide, linoleamide, linolenamide and arachidonamide, Fig. S1) also possessed this activity. These substances are produced by diverse bacterial^30^ and eukaryotic microbiota^31,32^ as well as plants^33–36^, and are even found in human serum^37^, tear film^38^ and breast milk^39^. We demonstrate that *cis*-unsaturated fatty amides fluidize Gram-negative bacterial membranes at micromolar concentrations, compromising the permeability barrier of the cell envelope. Furthermore, we show that oleamide restores the sensitivity of carbapenem-resistant *P. aeruginosa* to imipenem in a murine full-thickness wound model. Taken together, we propose that our culture-dependent platform and the associated screening techniques will have widespread use in identifying and understanding biotic factors that modulate antimicrobial susceptibility.

## RESULTS

### Many bacteria produce spent medium sensitizing P. aeruginosa to meropenem

A growing body of literature indicates that primary^40,41^ and secondary^42^ metabolites alter the susceptibility of bacteria to antibiotics. While most work has focused on metabolites leading to resistance^42^, less is known about the nature and prevalence of metabolites leading to enhanced antibiotic susceptibility. For example, some microbes co-produce multiple biomolecules that act synergistically to harm other microbial cells^43^, while others seem to produce *bona fide* antibiotic potentiators^26–29^. We therefore hypothesized that a variety of microbiota might produce soluble molecules that function to sensitize bacteria to antibiotics.

To test this hypothesis, we used medium-scale parallel culturing to amass a diverse, living library of bacteria and fungi. These organisms originated from human donors as well as animal, plant, aquatic, and soil specimens (see Methods, Supplementary Data 1). We varied agar nutrient compositions, oxygen levels and temperature to further isolate and select a subset of 348 strains – representing ≥157 species from 60 genera – in monocultures (Supplementary Data 2). All the strains selected were amenable to aerobic growth in brain-heart-infusion (BHI) broth. Subsequently, we devised a facile, medium-throughput assay to evaluate spent media from these organisms for the capacity to potentiate the activity of antibiotics against a target bacterium (Fig. 1A). The crux of the assay was calculating relative growth (see Methods, Eqn. 1). We define this value as the ratio of growth for the target organism in spent media containing an antibiotic versus the growth of that same organism in spent media alone. This calculation is based on measurements of optical density at 600 nm (OD_600_) normalized to internal controls in microplates (Eqn. 1). In principle, this approach provides an agnostic means to identify organisms producing spent medium that alters the susceptibility of a target bacterium to nearly any antimicrobial agent (Fig. 1A).

**Fig. 1.**
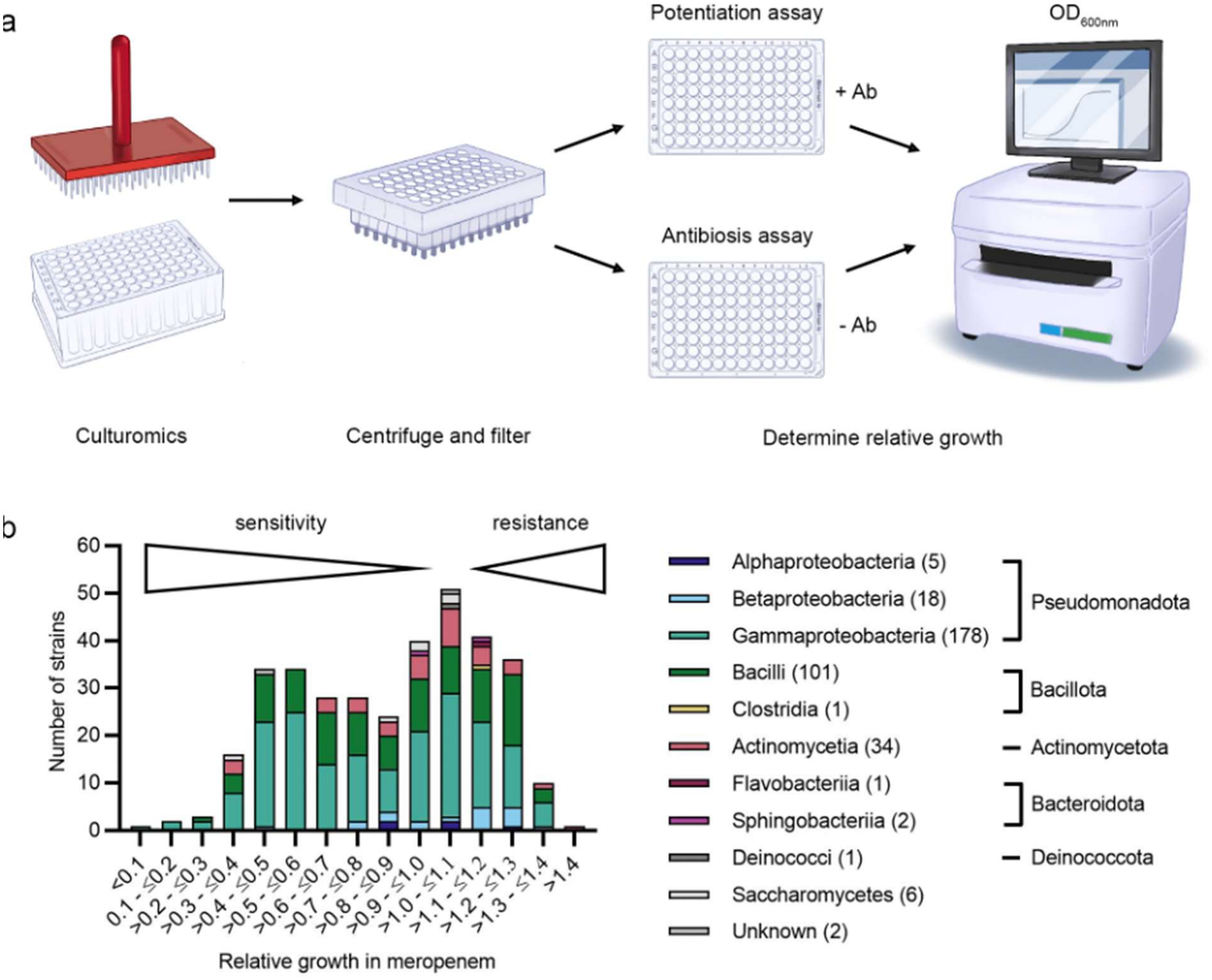
A culture-based, medium-throughput platform provides technology to identify microbial species producing spent media that potentiate last resort antibiotics against carbapenem-resistant P. aeruginosa. (a) Simplified workflow for a medium-throughput screen to identify spent media that can potentiate antibiotics. (Credit: Jesse Horne for original scientific illustration). (b) A frequency histogram revealing a skewed distribution in the number of strains producing spent media that potentiate meropenem against carbapenem-resistant P. aeruginosa M6A146. Bacterial and fungal phylogeny are color-coded (by class, phylum or division). Number of strains surveyed by culturomics in this study are indicated in parentheses.

Here, we strategically targeted a human isolate of carbapenem-resistant *P. aeruginosa* designated strain M6A146 (see Methods). This strain is multidrug resistant (Fig. S2). We observed that the minimum inhibitory concentration (MIC) of meropenem towards *P. aeruginosa* M6A146 was 64 µg/mL, which exceeds the clinical breakpoint of the drug (≤2 µg/mL)^44^. Analysis of the complete *P. aeruginosa* M6A146 genome revealed a nonsense mutation in the *oprD* gene linked to carbapenem resistance (see Methods).

In the screen, we sought to identify microbes producing spent media that resulted in inhibition of *P. aeruginosa* M6A146 in 2 µg/mL meropenem, a 32-fold decrease in meropenem resistance corresponding to the precipice of the clinical breakpoint. While we observed that spent medium from some strains increased the resistance of *P. aeruginosa* M6A146 to meropenem, a frequency histogram of mean relative growth values from this screen showed a distribution skewed towards values <1 (Fig. 1B). This non-normal distribution unexpectedly indicates that a disproportionate number of microbiota produced spent media increasing the susceptibility of *P. aeruginosa* M6A146 to meropenem (Fig. 1B).

We chose 20 bacterial strains producing spent media exhibiting the greatest decrease in relative growth (2.5- to 40-fold, Supplementary Data 2). We carried out secondary validation to control for false positives using viable cell counting at a bactericidal concentration of meropenem (32 µg/mL), which was otherwise sub-inhibitory to *P. aeruginosa* M6A146 (Fig. 2). This produced a high-confidence shortlist of 13 strains constituting candidate producers of substances sensitizing *P. aeruginosa* to meropenem (Fig. 2). Taken together, our data indicate that diverse bacterial species in pure monoculture produce spent media that potentiate meropenem activity against carbapenem-resistant *P. aeruginosa*, and moreover, that this capacity is remarkably prevalent.

**Fig. 2.**
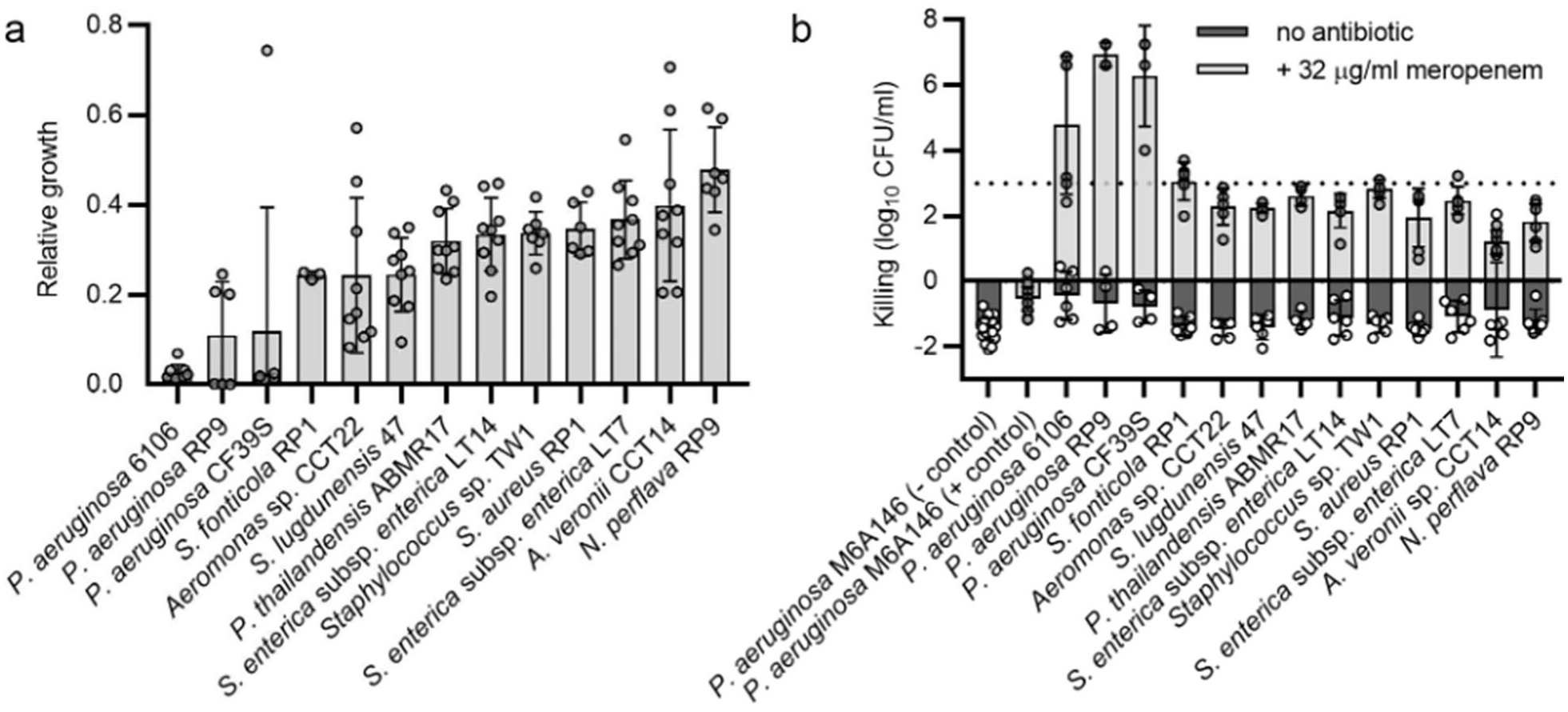
A medium-throughput screen identifies and validates that diverse bacterial species produce spent media that potentiate meropenem against carbapenem-resistant P. aeruginosa. (a) Relative growth calculation of P. aeruginosa M6A146 in fresh brain-heart infusion broth containing 2 µg/mL meropenem amended with spent media from the indicated bacteria. See text for details of the relative growth calculation, which is based on optical density measurements at 600 nm in deep-well microplates. (b) Log-killing of P. aeruginosa M6A146 in fresh brain-heart infusion broth containing 32 µg/mL meropenem amended with spent media from the indicated bacteria. In both panels, each datum point represents 1 to 3 technical replicates from 2 or 3 independent biological replicates. Bars and lines represent means and standard deviations, respectively.

### Oleamide is a natural antibiotic potentiator

The relative growth calculation corrects for changes in *P. aeruginosa* M6A146 yield that might depend on spent medium composition (Eqn. 1). It further allows us to identify and eliminate organisms producing antibiotics from the analysis. Additionally, meropenem has pKa_1_ and pKa_2_ values of 2.9 and 7.4, which correspond to its carboxylic acid and pyrrolidinylamino groups, respectively^45^. Changes in pH can affect not only the stability of meropenem in solution^46,47^, but also the *in vitro* susceptibility of *P. aeruginosa*^48^ and other bacteria^49^ to the antibiotic, especially at values ≤5.5 [refs. ^48,49^] and ≥9.0 [ref. ^48^]. However, pH testing of spent media that was combined with cation-adjusted Mueller Hinton broth (CA-MHB) used for screening showed a mean pH and standard deviation of 6.35 ± 0.43 for those strains exhibiting the greatest antibiotic potentiation (Fig. 2), which is marginal to the pKa_2_ value of meropenem. We thus sought other moieties in spent media that may account for the observed bioactivity.

We used a process of filtration, solvent extraction and fractionation to isolate bioactive matter (Fig. 3A). To begin, we focused on strains producing spent media containing small, bioactive moieties passing filtration with a molecular weight cut-off ≤10 kDa. We further narrowed in on a subset of these strains – *Neisseria perflava* RP9, *Staphylococcus lugdunensis* 47, and *Aeromonas* sp. CCT22 – all yielding bioactive moieties that partitioned into hydrophobic solvents during extraction steps.

**Fig. 3.**
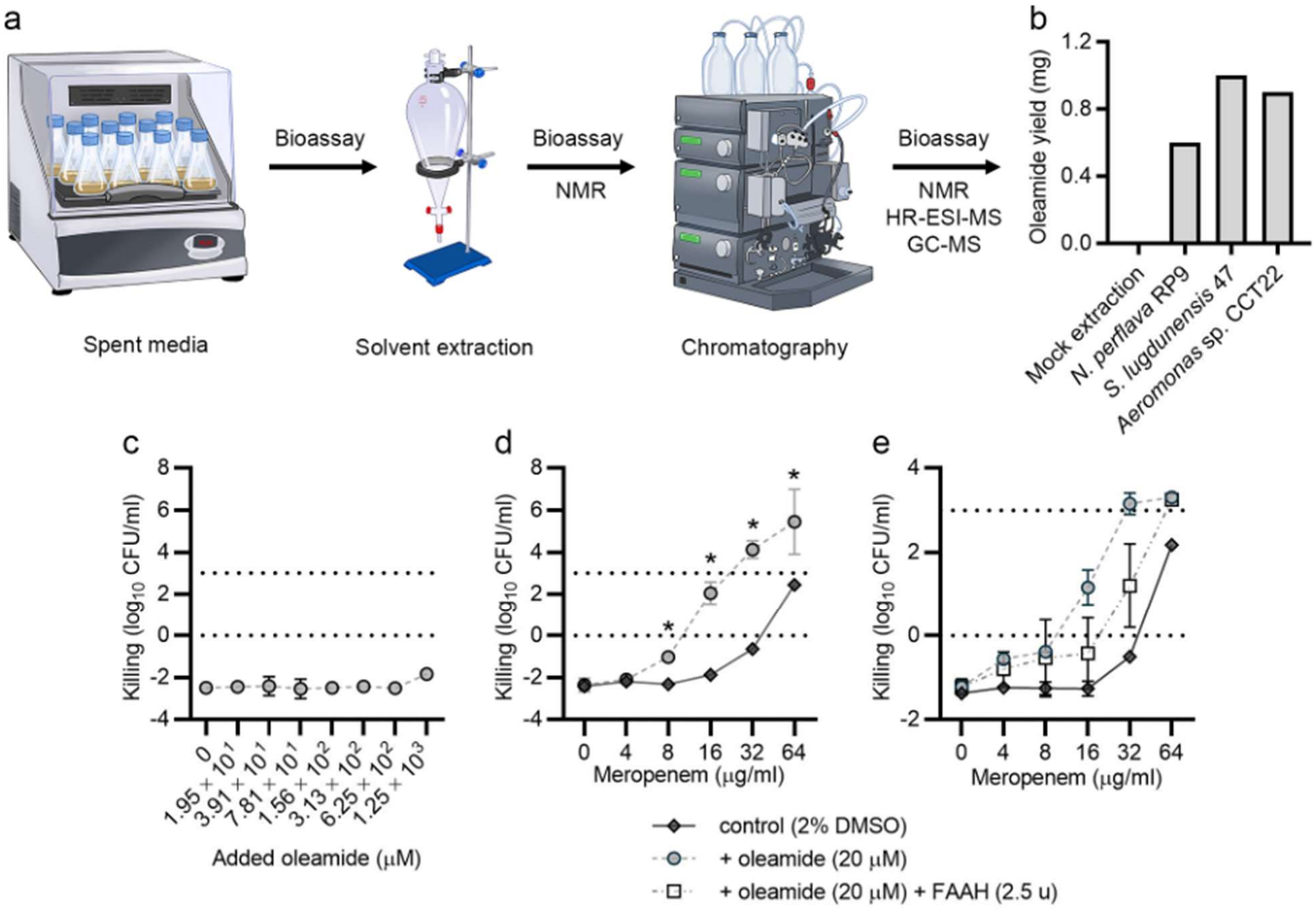
Bioassay-guided fractionation identifies that oleamide is a natural antibiotic potentiator produced by several bacterial species. (a) A simplified workflow for bioassay guided fractionation used in this study (Credit: Jesse Horne for original scientific illustration). (b) Yield of oleamide per liter of input culture from exemplar bioactive fractions. (c) Susceptibility or P. aeruginosa M6A146 to oleamide. (d) Susceptibility of P. aeruginosa M6A146 to meropenem with and without the addition of oleamide. (e) Reduction in meropenem potentiation by oleamide against P. aeruginosa M6A146 by fatty acid amide hydrolase (FAAH). In panels (c), (d) and (e), points and error bars represent the means and standard deviations of ≥3 independent biological replicates. Significant differences were calculated using a two-tailed Student’s t-test with the Holm-Sidak multiple test correction (* denotes p < 0.05).

It is difficult to predict candidate bioactive molecules from non-model organisms. We note that previous studies with *Staphylococcus aureus* led to the discovery that small, hydrophobic cis-unsaturated fatty acids sensitize some Gram-positive organisms to antibiotics^50–52^. However, this group of molecules is not known to display this activity against Gram-negatives^52^. Therefore, we used bioactivity-guided fractionation via chromatography, as required, to detect additional fractions of low chemical complexity exhibiting antibiotic potentiating activity (see Methods). Subsequently, nuclear magnetic resonance (NMR), high-resolution electrospray ionization mass spectrometry (HR-ESI-MS) and gas chromatography mass spectrometry (GC-MS) (Fig. 3A, see Methods) were used to identify molecules in these bioactive fractions. This process identified that all 3 of these species produced oleamide in their bioactive spent media fractions (Fig. 3B).

Although some previous work has attributed antimicrobial activity to oleamide^32,53,54^, we found that by itself, purified oleamide exhibited little if any antibacterial activity against *P. aeruginosa* M6A146 in these endpoint assays – even at added concentrations in excess of 1 mM (Fig. 3C). However, at a concentration as low as 20 µM and in combination with meropenem, oleamide reduced the MIC and minimum bactericidal concentration (MBC) of the antibiotic 4- to 8-fold (Fig. 3D), and increased *P. aeruginosa* M6A146 killing up to ∼13,000-fold at achievable bactericidal concentrations of the drug (Fig. 3D). We also found that oleamide potentiated imipenem and/or meropenem against many but not all other carbapenem-resistant *P. aeruginosa* clinical isolates^55^, including isolates expressing the Verona integron–encoded metallo-β-lactamase 7 (VIM-7) and the Guiana extended spectrum 5 (GES-5) carbapenemase, or known to possess loss-of-function mutations in *oprD* (Fig. S3). This spectrum of activity is consistent with knowledge that antibiotic resistance in *P. aeruginosa* is multifactorial and diverse from strain-to-strain^56^. We corroborated the antibiotic-potentiating activity of oleamide by adding recombinant human fatty acid amide hydrolase (FAAH) to media. FAAH is a serine hydrolase that hydrolyzes the amide bond of fatty amides like oleamide, yielding a fatty acid and a primary amine^57,58^. FAAH impeded meropenem potentiation by oleamide *in vitro* (Fig. 3E).

Oleamide is a lubricant used in plastics manufacturing^59,60^ and can leach from some types of disposable laboratory plasticware^59^. However, process controls for extractions contained no detectable oleamide (Fig. 3B). This result suggests that oleamide originates from the bacteria, even if the biosynthetic pathways in the non-model organisms *N. perflava* RP9, *S. lugdunensis* 47, and *Aeromonas* sp. CCT22 remain unknown. Also, we posit that there may be more than one bioactive moiety in spent media because purified oleamide did not reduce the *P. aeruginosa* M6A146 imipenem MIC to the same extent as spent media from these organisms; moreover, bioactivity was observed in multiple chemical fractions (not shown). Nevertheless, our data identify that oleamide is a natural antibiotic potentiator produced by a variety of microbiota, and that it exhibits bioactivity against different *P. aeruginosa* isolates.

### Chemically diverse cis-unsaturated fatty amides are antibiotic potentiators

Fatty amides are chemically related to fatty acids (Fig. S1). There is an extensive literature attributing antibacterial activity to fatty acids^61^, some of which may even enhance antibiotic activity^62^. However, we observed that the fatty acid, oleic acid, which lacks the amide group of oleamide (Fig. S1), exhibited no antibacterial or antibiotic-potentiating activity against *P. aeruginosa* M6A146 (Fig. S4). Thus, we investigated the chemical groups of oleamide and their arrangements that are linked to antibiotic potentiation using structure-activity relationship studies.

Here, we acquired fatty acids in which acyl chain length, orientation of double bonds, and degree of saturation were varied. We synthesized an array of corresponding fatty amides from them (see Methods). Subsequently, we ascertained the susceptibility of *P. aeruginosa* M6A146 to meropenem in the presence and absence of these fatty acids and fatty amides. Although twofold dilution gradients of meropenem were tested for all molecules (Figs. S4, S5, and S6), the results of this analysis were most evident at the bactericidal threshold of meropenem (32 µg/mL, Fig. 4A).

**Fig. 4.**
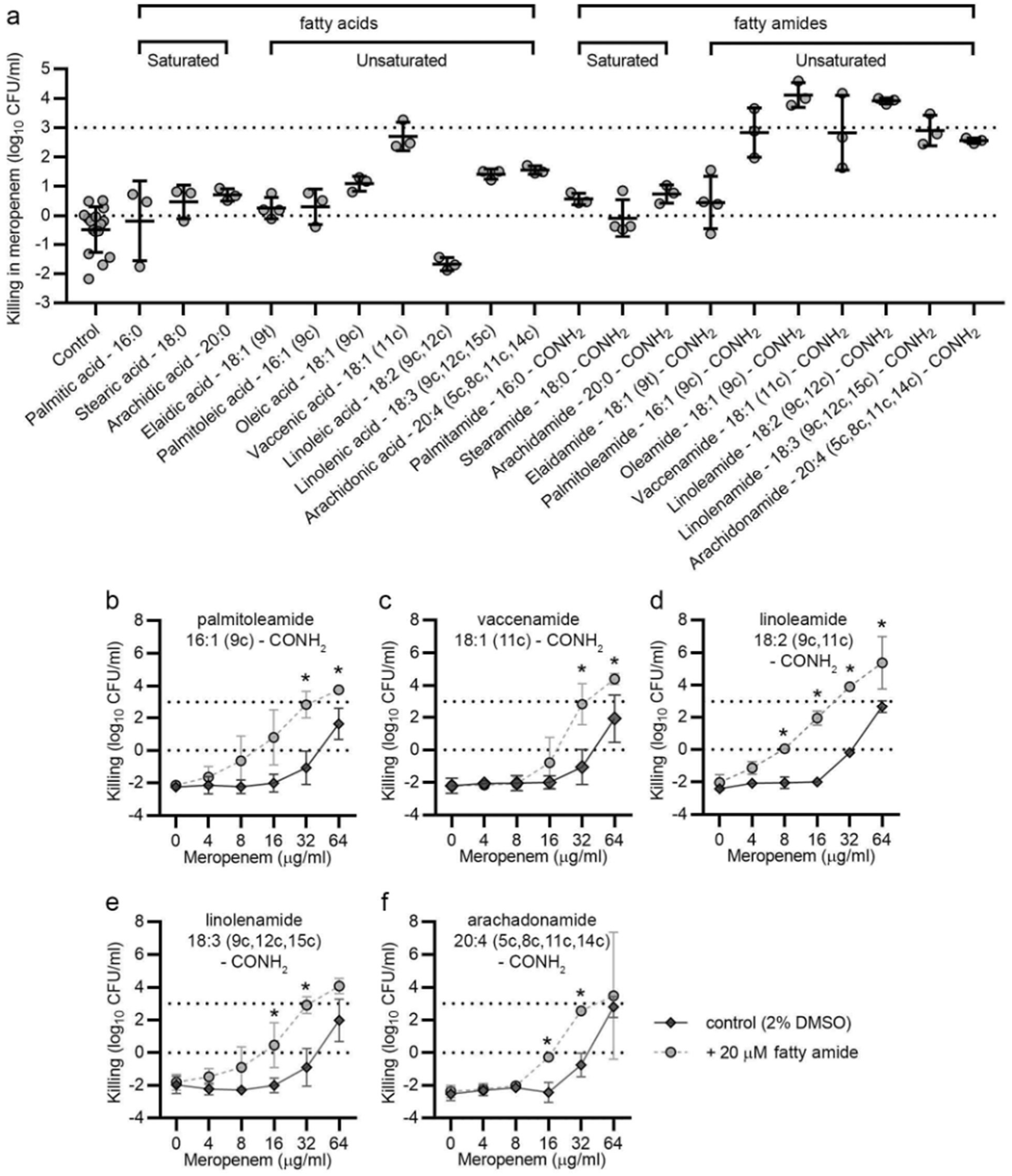
Structure activity relationship analysis of oleamide defines functional groups for a new class of antibiotic potentiators. (a) Susceptibility of P. aeruginosa M6A146 to fatty acids and fatty amides in meropenem. Datum points, lines and error bars represent single biological replicates, means, and standard deviations (SDs), respectively. Meropenem susceptibility of P. aeruginosa M6A146 in (b) palmitolamide, (b) vaccinamide, (c) linoleamide, d) linolenamide, and (e) arachadonamide. Points and error bars represent the means and SDs of ≥3 independent biological replicates. Significant differences were calculated using a two-tailed Student’s t-test with the Holm-Sidak multiple test correction(*denotes p<0.05).

**Fig. 5.**
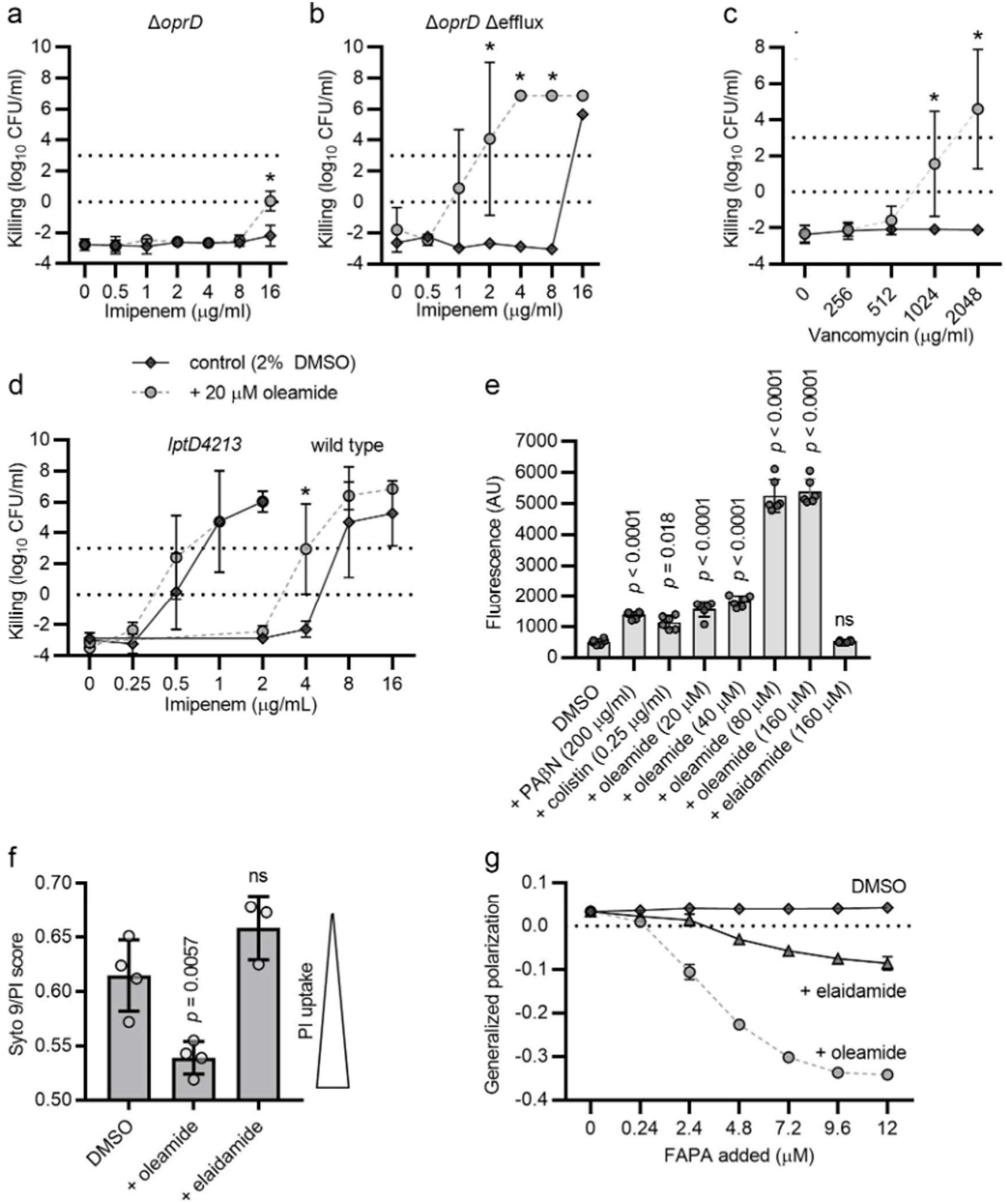
Oleamide holistically compromises the permeability barrier of the P. aeruginosa cell envelope, increasing small molecule uptake. Susceptibility of P. aeruginosa PAO1 (a) ΔoprD and (b) ΔoprD Δefflux mutants to imipenem with and without the addition of oleamide. (c) Susceptibility of P. aeruginosa M6A146 to vancomycin with and without the addition of oleamide. (d) Susceptibility of P. aeruginosa PAO1 wild type and mutant strains to imipenem with and without the addition of oleamide. Points and error bars represent the means and standard deviations (SDs) of ≥3 independent biological replicates. Significant differences were calculated using a two-tailed Student’s t-test with the Holm-Sidak multiple test correction (* denotes p < 0.05). (e) Ethidium bromide uptake by P. aeruginosa M6A146. P values calculated using Student’s t-test to compare test groups vs. the control with 2% DMSO. Circles represent biological replicates; bars and lines represent means and SDs. P values were calculated using a two-tailed Student’s t-test. (f) Propidium iodide uptake by P. aeruginosa M6A146 using ImageJ to quantify green and red pixels in fluorescent microscopy images (see Methods). Circles represent mean values from a single field of view for a biological replicate; bars and lines represent means and SDs. P values were calculated using a two-tailed Student’s t-test. (g) Generalized polarization of large unilamellar vesicles designed to model the P. aeruginosa inner membrane using laurdan GP (see Methods). Symbols and bars represent means and SDs of 3 independent replicates.

Under the conditions tested, none of the fatty acids or fatty amides had intrinsic antibacterial activity alone (Figs. 4, S4, S5 and S6). Additionally, the saturated fatty acids and fatty amides exhibited no capacity for antibiotic potentiation (Figs. 4 and S4). Meropenem combined with some *cis*-unsaturated fatty acids (vaccenic, linoleic, linolenic and arachidonic acids) displayed low but statistically significant increases in killing over very narrow concentration ranges (Fig. 4 and S5). By contrast, all tested *cis*-unsaturated fatty amides potentiated meropenem over a broader range of concentrations (Fig. 4B – F), providing, maximally, ∼10,000-fold decreases in viable cell counts relative to untreated controls (Fig. 4B – F). Although fatty amides with C_16_, C_18_ and C_20_ acyl chains all exhibited bioactivity (Fig. 4), the most active fatty amides had C_18_-acyl chains with 1 or 2 *cis* double bonds, with one of the double bonds at the C_9_ position (oleamide and linoleamide, Fig. 3C and 4D, respectively). The *cis* orientation of double bonds was essential. Fatty acids and fatty amides with *trans* double bonds (elaidic acid and elaidamide, respectively, Fig. S1) displayed no bioactivity (Fig. 4A and Fig. S6). Taken together, these data reveal that other fatty amides – most of which also occur naturally – potentiate antibiotics. Moreover, structure-activity relationship studies indicate that the amide and *cis*-oriented double bonds are key functional groups for the bioactivity of these molecules.

### Oleamide does not inhibit β-lactamases

We next sought to understand how fatty amides exert antibiotic-potentiating activity. Here, we turned investigations towards the laboratory *P. aeruginosa* PAO1 strain because of its genetic tractability^63–65^, providing a straightforward way to use engineered genotypes (Table S1 and S2) to identify, validate, or eliminate putative cellular targets from the analyses.

Early clues about mode-of-action resulted from investigating the ability of oleamide to potentiate other classes of antibiotics. A curious observation was that oleamide potentiated imipenem against *P. aeruginosa* PAO1, but not other β-lactams representing the cephalosporin or monobactam antibiotic classes (Fig. S7). The targets of all β-lactam antibiotics are the penicillin binding proteins involved in peptidoglycan synthesis and localized to the periplasmic space^66^. However, perhaps with the notable exception of the carbapenems, many antibiotics, including most β-lactams, do not appear to require porins to bypass the outer membrane^67^. Resistance to these other β-lactams likely depends more on other cellular factors, including the β-lactamases^68^. We thus hypothesized that fatty amides must not target β-lactamases.

To test this hypothesis, we measured β-lactamase activity in *P. aeruginosa* M6A146 periplasmic fractions with and without exogenous oleamide. We found that oleamide had no significant effect on β-lactamase activity (Fig. S8A). Furthermore, oleamide did not potentiate cefotaxime activity against *P. aeruginosa* PAO1 that was engineered for inducible overexpression of the *ampC* gene (Fig. S8B), which encodes a β-lactamase that can degrade cephalosporins like cefotaxime^69^. Taken together, these data suggest that fatty amides do not inhibit or target β-lactamases when potentiating carbapenems against *P. aeruginosa*.

### Active efflux conceals a role for oleamide in facilitating antibiotic uptake

We further probed the ability of oleamide to potentiate imipenem against a *P. aeruginosa* PAO1 Δ*oprD* mutant, which, analogously to *P. aeruginosa* M6A146, has a loss-of-function mutation in the gene encoding the OprD carbapenem-uptake porin. We noted a ≥4-fold increase in the imipenem MIC of the Δ*oprD* mutant relative to the wild type strain (Fig. 5). Oleamide potentiated imipenem against the PAO1 Δ*oprD* mutant; however, the magnitude of sensitization was limited (Fig. 5A). Because *P. aeruginosa* possesses several trans-envelope efflux pumps that can extrude antibiotics^7–9^, we hypothesized that active antibiotic efflux might reduce the observable antibiotic-potentiating activity of oleamide. To test this hypothesis, we examined imipenem potentiation in an isogenic PAO1 Δ*oprD* mutant that also lacked 15 other genes encoding RND-type multidrug efflux pumps (*mexAB*-*oprM*, *mexCD*-*oprJ*, *mexXY*, *mexJKL*, *mexHI*-*opmT*, and *opmH*), named PAO1 Δ*oprD* Δefflux^55,70^. We saw striking oleamide-dependent potentiation of imipenem in the Δ*oprD* Δefflux background (Fig. 5B), observing up to 1,000,000-fold increases in killing over a 16-fold range of imipenem concentrations relative to untreated controls (Fig. 5B). Although oleamide did not potentiate meropenem against *P. aeruginosa* PAO1 (Fig. S7) – and the meropenem MIC of the Δ*oprD* mutant (2 µg/mL, Fig. S9A) was unchanged relative to the wild type strain (Fig. S7) – we observed that oleamide provided similarly striking potentiation of meropenem against the Δ*oprD* Δefflux strain (Fig. S9B). Taken together, these data suggest that efflux activities across membranes are not primary targets of oleamide; rather, oleamide potentiates imipenem by facilitating antibiotic uptake.

### Oleamide holistically compromises the permeability barrier of the cell envelope

Another key observation was that oleamide potentiated vancomycin against *P. aeruginosa* PAO1 (Fig. S10) and M6A146 (Fig. 5C), albeit at high concentrations. The vancomycin glycopeptide exhibits a Gram-positive spectrum of activity, owing to its inability to permeate the Gram-negative outer membrane^12,14^. Alongside the data that oleamide potentiates outer membrane-impermeable imipenem against *P. aeruginosa* PAO1 (Fig. S7), this hypersensitivity suggests that oleamide facilitates antibiotic transit across the outer membrane. This putative mode-of-action was queried by measuring the imipenem sensitivity of a *P. aeruginosa* PAO1 strain with a mutation in LptD, the final protein of LPS synthesis and transport to the outer membrane. This mutant bears an *lptD4213* allele that compromises outer membrane barrier function^5^. We hypothesized that if oleamide targeted an outer membrane function already missing due to the *lptD4213* mutation, then no further potentiation would occur. Indeed, we observed that oleamide did not potentiate imipenem against the *lptD4213* mutant (Fig. 5D), providing additional evidence that oleamide facilitates small molecule transit across the outer membrane.

We also found that oleamide potentiated representative aminoglycoside, fluoroquinolone, tetracycline and polymyxin antibiotics (Fig. S10). With the exception of polymyxin, all have cytosolic targets. These data imply that oleamide may alter permeability of the inner membrane too. Here, we directly assessed small molecule accumulation in the cytoplasm of oleamide-treated cells by measuring cellular accretion of ethidium bromide (EtBr), a nucleic acid intercalator that fluoresces on binding cytosolic DNA and RNA. Oleamide treatment increased EtBr accumulation in *P. aeruginosa* cells in a fashion similar to that exhibited by colistin and phenylalanine-arginine β-naphthylamide (PAβN), known chemical perturbators of membranes and/or xenobiotic efflux^71^. By contrast, elaidamide, the trans-unsaturated isomer of oleamide (Fig. S1), did not exhibit this activity (Fig. 5E).

We also assessed inner membrane integrity by imaging bacteria stained with Syto-9 and propidium iodide, the latter of which is thought only to enter the cytosol of cells with a compromised inner membrane^72^. Unsurprisingly, oleamide increased propidium iodide staining of cells, whereas elaidamide did not (Fig. 5F). These cells were grown under identical conditions used for susceptibility testing in which there were no significant differences in viable cell counts between oleamide and elaidamide-treated bacteria and solvent controls (Fig. 3 and S6). Thus, this result reveals a lack of correlation between propidium iodide staining and cell viability. We posit that this observation is consistent with the function of a *bona fide* antibiotic potentiator, which might exert little toxicity alone while otherwise augmenting the uptake of another molecule.

Because structure-activity relationship studies indicated that *cis*-oriented double bonds are essential for the bioactivity of fatty amides, we hypothesized that the shape of the oleamide molecule might disrupt the lipid packing and fluidity of membranes. Here, we sought to model the interactions of oleamide with the inner membrane using large unilamellar vesicles comprised of 1-palmitoyl-2-oleoyl-sn-glycero-3-phosphoethanolamine (POPE) and 1-palmitoyl-2-oleoyl-sn-glycero-3-phospho-(1′-rac-glycerol) (POPG, see Methods)^73^, mimicking the two most abundant phospholipids, phosphatidylethanolamine and phosphatidylglycerol, found in *P. aeruginosa*^74^. We tested the effect of oleamide on membrane fluidity by preparing large unilamellar vesicles with an embedded dye, Laurdan GP, which has an H_2_O-sensitive fluorescence emission spectrum^73^. Because the penetration of H_2_O into hydrophobic membranes depends on polar lipid head group packing density and fatty acid disorder, the emission spectrum of membrane-embedded Laurdan GP responds to membrane fluidity^73^. Indeed, we observed that oleamide significantly reduced the generalized polarization of membranes relative to solvent controls, and to a significantly greater extent than its trans-unsaturated isomer, elaidamide (Fig. 5G). Taken together, these data suggest that antibiotic-potentiating *cis*-unsaturated fatty amides affect lipid packing and membrane fluidity, and holistically disrupt the permeability barrier of the *P. aeruginosa* cell envelope.

### Oleamide potentiates the lactoferricin antimicrobial peptide

Oleamide is found in mammalian serum^37^, tear film^38^, and breast milk^39^. There is increasing recognition that host lipids may have antimicrobial functions^75,76^, and therefore, we theorized that oleamide might act to potentiate other antimicrobial components of innate immunity. A well studied antimicrobial protein is lactoferrin^77^, which is abundant in many bodily fluids – including serum^77^, tears^78^, and breast milk^77^. Various antimicrobial peptides are released from the proteolytic degradation of lactoferrin^79^, such as lactoferricin^79^. Lactoferricin is thought to interact with the polyanionic outer membrane surface of diderms to disrupt membrane fluidity thereby allowing the peptide to self-translocate into the cell to act against additional intracellular targets^79^. We reasoned that oleamide-linked changes in membrane lipid packing would favor the self-promoted uptake of lactoferricin; so, we measured the susceptibility of *P. aeruginosa* PAO1 to lactoferricin B in the presence and absence of oleamide. We observed that oleamide decreased bacterial viability >10,000,000-fold at a lactoferricin B concentration otherwise bacteriostatic to *P. aeruginosa* (Fig. S11). Our results demonstrate that, at least *in vitro*, oleamide significantly augments the activity of an antimicrobial peptide found in innate host defense.

### Oleamide augments the efficacy of topical antibiotics in vivo

Fatty amides are not ideal as systemic therapeutics because of their low water solubility and natural metabolism in mammals. However, analogous to host-derived unsaturated fatty acids that sensitize *S. aureus* to aminoglycosides *in vitro*^50,51^ and *in vivo*^52^, we posited that combinations of *cis*-unsaturated fatty amides with antibiotics might be useful topicals. We hypothesized that a composition of imipenem with oleamide – a fatty amide thought to pose low potential health risk and classified as a medicinal natural health product by Health Canada – could be used in wounds against carbapenem-resistant *P. aeruginosa* exhibiting resistance due to reduced drug uptake.

We tested this hypothesis using an established, full-thickness wound model in outbred, immunocompetent, SKH-1 hairless mice^52^ (Fig. 6A). To demonstrate that mode-of-action *in vivo* was due to overcoming reduced drug uptake, wounds were infected with the carbapenem-resistant *P. aeruginosa* PAO1 *ΔoprD* Δefflux strain. We observed that infected wound tissue treated twice daily with imipenem or oleamide alone showed no significant difference in bacterial load (Fig. 6B). By contrast, animals treated with imipenem and oleamide in combination had significantly fewer bacteria in wound tissue (∼16-fold less, Fig. 6B). We noted no significant differences in the number of *P. aeruginosa* in the spleen or kidneys between treatment and control groups (Fig. 6C and D). Taken together, we conclude that fatty amides may be used to augment antibiotic efficacy *in vivo*, and that our screening approach can yield natural products to inspire the design of experimental therapeutics.

**Fig. 6.**
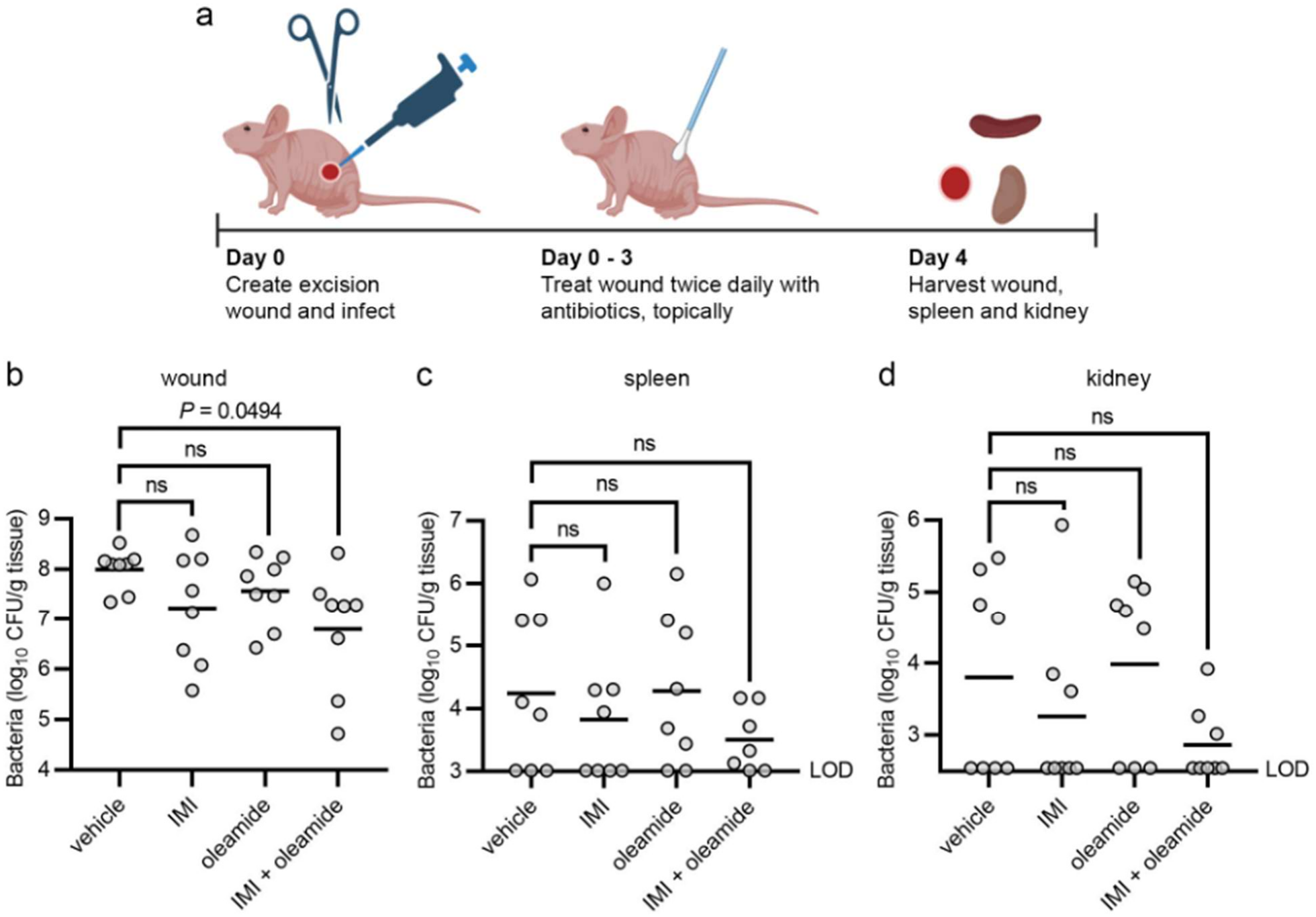
Oleamide potentiates imipenem against carbapenem-resistant P. aeruginosa in mouse wounds. (a) Schematic representing the wound infection model. A 4-mm full-thickness circular wound was created on the back of 8-10 week-old SKH-1 outbred immunocompetent mice. The wound was infected with ∼6 × 10^7^ CFU of P. aeruginosa MB6476 (ΔoprD Δefflux). Wounds were treated topically with the vehicle, 3% oleamide, 8 µg/ml imipenem, or both, twice daily commencing 1 h post-infection. On day 4 post-infection, mice were euthanized, and the bacterial burden was enumerated in (b) wound, (c) spleen, and (d) kidneys. Statistical significance was determined using a Brown-Forsythe and Welsh ANOVA multiple comparison test (n = 8 mice/group). LOD, limit of detection; ns, not significant.

## DISCUSSION

We present a culture-based, microplate compatible microbial natural products platform for identifying antibiotic potentiators. Future potential to vary culture conditions (e.g., O_2_, nutrients, etc.), use any microbial strain collection, and automate workflows provides for a highly versatile hit-to-lead platform to identify microbial species producing previously uncharacterized natural substances that potentiate antibiotics and other antimicrobials.

As proof-of-principle, we identified that several bacterial species produce oleamide (Fig. 3), which potentiates many antibiotics, including last resort carbapenems, against *P. aeruginosa*. Chemically related fatty amides exhibited analogous bioactivity (Fig. 4). Antibacterial activity has been reported for some fatty amides, including oleamide, which are sometimes referred to as fatty acid amides or *N*-acylamides^32,53,54^. By contrast to the primary fatty amides examined in this study, these antibacterial fatty amides more notably encompass secondary and tertiary fatty amides with synthetically derivatized amide groups^80,81^. Such molecules include derivatives of the antifungal, cerulenin^82^. However, antibiotic potentiation is an activity previously unreported among all these fatty amides, including the primary fatty amides. Primary fatty amides are prevalent in nature and are produced by organisms from all kingdoms of life, including diverse microbiota.

Oleamide is an endogenous signaling lipid. Oleamide and enzymes involved in its homeostasis, such as FAAH, have intricate roles in plant and animal physiology^58,83^, including inflammation^84^ and response to infection^85,86^. It is tempting to speculate, therefore, that a natural role for cis-unsaturated primary fatty amides might be to sensitize pathogens to antimicrobial molecules found in innate host defence, including peptides that disrupt bacterial membranes to translocate into bacterial cells, such as lactoferricin.

Antibiotic potentiators provide hope for clinicians that are struggling to treat multidrug-resistant bacterial infections. They are an important avenue to rescue older antibiotics and build capacity as new mechanisms of antimicrobial resistance emerge in clinics. Here, the natural metabolism and multifunctionality of fatty amides *in vivo* limits the design of antimicrobial compositions containing them, especially for the treatment of systemic infection. Optimizing pharmacokinetics and pharmacodynamics for co-administered molecules is a considerable challenge in drug development too. Nevertheless, our proof-of-concept data from a mouse model of full-thickness wounds (Fig. 6) indicates that it may be possible to further develop topical antimicrobial compositions containing fatty amides and antibiotics to treat drug resistant bacteria in wound infections.

*In vitro* evidence suggests that some antibiotic potentiators – such as the β-lactamase inhibitor tazobactam – may select for antimicrobial resistance^87^. Combination therapies are not foolproof, and we do not know how fatty amides may select for resistance. However, a theme from a variety of experimental models is that two or more antimicrobials in combination should delay the emergence of resistance^88–92^. For example, this could occur through a requirement for multiple resistance-linked mutations^88,93^, synergistic killing that reduces the population size^91^, or reducing the range of drug concentrations at which selection can act^94^. Clinical evidence supporting the administration of multiple antibiotics simultaneously to prevent antibiotic resistance development in humans remains a knowledge gap^95^.

While further studies are required to understand the potential toxicity of fatty amides used in topical antimicrobial formulations, several primary fatty amides, including oleamide, are thought to be of low risk to human health. For example, oleamide is ingested in small quantities daily by many people. Oleamide can leach from some food packaging and containers^96^. It is present naturally in some foods, including dairy products fermented with *Penicillium candidum*, such as Camembert cheese^97^. Isopropyl and diethyl derivatives of oleamide and linoleamide are widely used in cosmetics and personal care products and are considered safe when formulated to be non-irritating^98,99^.

Bacteria in diverse communities display a variety of competitive behaviours^100^. However, it is challenging to infer the significance of our data to microbial ecology. It is conspicuous that many of the bacteria producing spent media eliciting enhanced antibiotic activity against *P. aeruginosa* are other Gammaproteobacteria (Fig. 1B). In fact, other strains of *P. aeruginosa* produced spent media leading to the greatest decreases in meropenem resistance of *P. aeruginosa* M6A146, perhaps reflecting a predilection of *P. aeruginosa* to engage in intraspecific competition^101–103^. However, the ecological functions of natural antibiotic potentiators remain unknown and may be tricky to decipher. Some analogous questions still surround the natural functions of antibiotics^104–106^. Here, key barriers for studying fatty amides in ecology are the lack of knowledge about fatty amide biosynthesis and the absence of genetic systems for non-model microorganisms producing fatty amides.

Our limited knowledge of the chemical properties required for Gram-negative outer membrane penetration makes it difficult to design molecules with improved permeability. Moreover, designer potentiators bypassing the outer membrane often do not consider the added necessity of penetrating the inner membrane, which selects for permeability properties (hydrophilicity) orthogonal to that of the outer membrane (hydrophobicity). The medium-scale parallel culture-based platform described here constitutes a discovery pipeline that can identify natural molecules overcoming this problem. Hydrophobic molecules functioning in this manner are likely unrepresented in small molecule libraries as their abilities to partition into an aqueous phase, as ascertained using partition coefficients (e.g., logP values), contradict the typical ligand efficiency metrics used in medicinal chemistry^107,108^.

## METHODS

### Standard microbiological and molecular biology methods

All previously published and genetically engineered laboratory bacterial strains are listed in Table S1. All plasmids and primers used in genetic manipulations are listed in Table S2 and Table S3, respectively. All basic microbiological and molecular procedures were executed according to standard protocols^109^. Except where noted otherwise, DNA concentrations were measured via the A_260_/A_280_ method using a Nanodrop 2000 Spectrophotometer (ThermoFisher Scientific). Protein concentrations were determined using the Pierce 660 nm Protein Assay Reagent (ThermoFIsher Scientific), which was calibrated using bovine serum albumin standards. Genomic DNA (gDNA) isolation, plasmid preparation and DNA gel extraction were performed using nucleotide purification kits purchased from Qiagen or BioBasics. Taq DNA polymerase was purchased from New England Biolabs. Phusion DNA polymerase and BP Clonase II were purchased from ThermoFisher Scientific. Transformations of *P. aeruginosa* were carried out using established protocols for electroporation^65^. Oligonucleotide primers were purchased from Integrated DNA Technologies. Sterile filtration of buffers and media was executed as required using asymmetrical polyethersulfone (aPES) filters with a 0.2 µm pores.

### Ultrapure water and buffers

All ultrapure water used for buffers and media was prepared in-house using a Milli-Q Direct Water Purification System (Millipore-Sigma). Phosphate buffered saline (PBS) was purchased as a 20 × concentrate (Amresco). It was diluted as needed in ultrapure water that was sterilized using an autoclave. A 300.0 mM sucrose solution, which was used for electroporation, was made using 102.7 g sucrose brought up to a volume of 1.0 L, which was filter sterilized.

### Microbiological media for routine culture

*P. aeruginosa* and *E. coli* strains were routinely propagated in lysogeny broth (LB). LB was made with 10 g tryptone, 5.0 g yeast extract and 5.0 g NaCl per 1.0 L of ultrapure water. No salt LB (NSLB) + 15% w/v sucrose was made by autoclaving 700.0 mL of ultrapure water containing 10 g tryptone and 5.0 g yeast extract prior to the addition of 300.0 mL of a 50% w/v sucrose solution, the latter of which was prepared by dissolving 500.0 g sucrose in 1.0 L ultrapure water and filter sterilized. Brain-heart infusion (BHI) broth and cation-adjusted Mueller-Hinton broth (CA-MHB) were purchased from BD Difco and prepared according to manufacturer’s directions. Semi-solid agar in Petri dishes was prepared by adding 15.0 g/L bacteriological agar to LB, NSLB or CA-MHB-based media prior to autoclaving. All strains were stored at −70 °C in LB containing 16.7% glycerol.

Vogel-Bonner Minimal Medium (VBMM)^110^ was prepared as a 10 × stock containing 2.0 g MgSO_4_•7H_2_O, 20.0 g citric acid, 100 g K_2_HPO_4_, and 35.0 g NaNH_4_HPO_4_•4H_2_O that was adjusted to pH 7.0 and filter sterilized. VBMM was prepared, as required, by diluting this stock 10-fold with autoclaved ultrapure water, and by adding 1.0 mL of a 1000 × trace metals solution^111^ to it. The 1000 × trace metal solution contained 13.5 x 10^-2^ g FeCl_3_•6H_2_O, 494.8 x 10^-3^ g MnCl_2_•4H_2_O, 3.7 x 10^-3^ g Zn(NO_3_)_3_•6H_2_O, 27.7 x 10^-3^ g CaCl_2_, 7.0 x 10^-3^ g CoSO_4_•7H_2_O, 4.3 x 10^-3^ g CuSO_4_, 6.0 x 10^-3^ g NaMoO_4_•H_2_O, 123.2 x 10^-3^ g MgSO_4_•7H_2_O and 32.4 x 10^-3^ g NiCl_2_ per 250.0 mL of ultrapure water, which was filter sterilized and stored at −20 °C. VBMM agar was made by adding 100 mL 10 × VBMM and 1 mL of the 1000 × trace metal solution to 900 mL of ultrapure water containing 10 g noble agar that had been autoclaved.

### Antibiotics

Aztreonam (ATM), carbenicillin (CAR), cefotaxime (CTX), colistin (CST), ciprofloxacin (CIP), imipenem (IPM), meropenem (MEM), tetracycline (TET), tobramycin (TOB), and vancomycin (VAN) were purchased from Sigma-Aldrich. Gentamicin (GEN) was purchased from Bio Basic. All stock solutions were corrected for activity, filter sterilized, split into 0.5 mL aliquots, and stored at −70 °C until used. Antibiotics were added as follows to media for routine culture and selection during genetic manipulations: for *E. coli*, CAR at 10 µg/mL, and GEN at 10 µg/mL; for *P. aeruginosa* PAO1: GEN at 60 µg/mL for two-step allelic exchange and 30 µg/mL for miniTn7 mutagenesis; for *P. aeruginosa* M6A146: MEM at 15 µg/mL.

### Viable cell counts

Where necessary, viable cell counts were ascertained by established protocols^112^ using serial tenfold dilutions of 20 µl aliquots of cultures in microplates, and by spotting 5 µl aliquots of these serially-diluted cultures on LB agar. These agar plates were incubated overnight at 30 °C before colony growth was scored.

### Medium-scale parallel culturing and strain isolation

We used a modified approach to culturomics (a process that seeks to exhaustively isolate microbiota from a target ecosystem using multiple culture conditions and mass spectrometry and/or 16S rRNA gene sequencing for isolate identification^113^) to enable the rapid construction of a diverse, living library of bacteria that could be probed for bioactive substances. A challenge in culturomics is redundancy in the screening, requiring the isolation and identification of up to hundreds of thousands of colonies from agar media to recover a few hundred species^113^. Here, we reasoned that shallow sampling of numerous, dissimilar environments would capture the greatest breadth in microbial biodiversity, while significantly reducing the number of colonies from agar that need to be isolated and identified.

Various dehydrated microbiological media and supplements were obtained from suppliers and prepared according to standard methods^114–117^ (formulations appear in Table S4 and S5). We cultured microbiota from a variety of environmental samples and human volunteers using 34 different types of agar media, varying oxygen levels and incubation temperatures. The isolation medium, atmospheric conditions, and incubation temperature for every strain isolated and used in the screen as well as salient contextual data appear in Supplementary Data 1. All of these strains were reposited in the Alberta Microbiota Repository (AMBR, see Data Availability).

### Human donors, animal models and research ethics

Human microbial isolates investigated were isolated from anonymous, secondary biological materials from healthy individuals. As anonymous biological samples were used to isolate microorganisms, formal ethics review was not required in accordance with policies of 1) the Conjoint Faculties Research Ethics Board (CFREB) at the University of Calgary, and 2) the Canadian Tri-Council Policy Statement 2 (2022) for the ethical conduct of research involving humans. All animal protocols were approved by the Institutional Animal Care and Use Committee at the University of North Carolina at Chapel Hill and met guidelines of the US National Institutes of Health for the humane care of animals.

### Microbial isolate data standardization and interoperability

To capture contextual data (e.g., sample context, growth and isolation conditions, sequencing techniques, bioinformatic processes, etc.) for strains isolated from massively parallel culturing, we developed a bespoke data specification reusing existing, open-source, community-developed standards and ontologies that was implemented in the DataHarmonizer curation tool^118^. DataHarmonizer provides an easy-to-use, spreadsheet-like template to users, as well as curation features and data validation functionality. The template is supported by a curation standard operating procedure as well as field and term reference guides. The specification is open source, version controlled, updated according to evolving data needs, and publicly available (see Data Availability).

### Identification of bacterial and fungal isolates by 16s or ITS rRNA gene sequencing

Bacterial 16s rRNA genes were amplified using a colony PCR protocol with primer pairs 27F/926R or 27F/1492R (Table S3) as required. ITS DNA regions were amplified using the primer pair ITS1-F/ITS4 (Table S3). If colony PCR did not yield a product detectable by analytical gel electrophoresis, then gDNA was extracted from bacteria or fungi grown in monoculture on the appropriate agar medium, and PCR was repeated with purified gDNA. In all cases, PCR products were purified from reaction mixtures and sent for Sanger sequencing with the primers used to target the amplicon.

Forward and reverse reads were manually trimmed for quality (i.e. only unambiguous base-calls from chromatograms were included). The bidirectional reads were then assembled using the Geneious^119^ Prime (2021) De Novo Assemble algorithm with default settings (highest sensitivity). Assembled 16s RNA gene sequences were queried against the curated EZBioCloud database^120^ of 16s rRNA gene sequences (version 2021.07.07). For bacteria, we made presumptive genus-level identifications based on a query sequence that had either 1) >98.7% identity to a type (T) sequence and/or >99.0% identity with a sequence that had not been typed, or 2) >99.0% identity with >1 type (T) sequence in the database. We made presumptive species-level identifications based on a query sequence with >99.3% identity to a type sequence and only if that match was unambiguous (i.e. there was >99.3% identity with only 1 type sequence in the database). Species designations were made using reference sequences from the List of Prokaryotic names with Standing in Nomenclature (LPSN)^121^. By contrast, assembled internal transcribed spacer (ITS) sequences were queried using BLASTN to search the National Center for Biotechnology Information (NCBI) ITS database. We made presumptive genus and species-level identifications based on expect (e) scores. 16s rRNA and ITS gene sequences for all organisms identified in this study appear in Supplementary Data 1.

### *P. aeruginosa* M6A146 genome sequencing, assembly, and annotation

gDNA was purified from 2.0 mL of an overnight culture of *P. aeruginosa* M6A146 using the DNeasy Blood and Tissue Kit (QIAGEN). The manufacturer’s protocol was followed except that gDNA was treated twice with RNAse A (ThermoFisher Scientific) and the final elution step was carried out with 100 µl of 10 mM Tris-HCl buffer (pH 8.0). DNA concentrations were measured using the Qubit v2.0 dsDNA HS Assay Kit (ThermoFisher Scientific). The purified *P. aeruginosa* M6A146 gDNA was sent to Genome Quebec for sequencing on the Pacific Biosciences Sequel platform. Briefly, 2.0 μg of high molecular weight gDNA was sheared using the Covaris g-TUBES (Covaris Inc., Woburn, Massachusetts, USA) at 1750 × g for 60 s in Centrifuge 5424 (Eppendorf).The gDNA libraries, which had an average insert size of 7 kb or larger, were prepared using the SMRTbell Express Template Prep Kit 2.0 (Pacific Biosciences). Sequencing primer v4 was annealed at final concentration of 1 nM and the Sequel 3.0 polymerase was bound at 0.5 nM. The libraries were purified using the AMPure XP Reagent (Beckman Coulter) according to the SMRTlink v7 calculator procedure. The DNA libraries were then sequenced on a PacBio Sequel instrument at an on-plate loading concentration of 5 pM using the diffusion loading protocol, using the Sequel Sequencing Plate 3.0, SMRT cells 1M v3 and 10 h movies.

Assembly of a single 6,755,766 bp contig – produced from 69,570 filtered subreads with a mean length of 7,932 bp and providing 87.6-fold coverage of the genome – was carried out by the Canadian Center for Computational Genomics (C^3^G) using a Hierarchical Genome Assembly Process (HGAP) workflow^122^ in the SMRT Analysis v2.3.0.140936.p5 software suite (PacBio). The subreads were filtered based on their length and QV score in order to identify the shorter, highly accurate reads as well as the longer, less accurate ones. An initial pre-assembly was carried out by aligning the high-quality short reads to the long reads with BLASR. The consensus sequences were generated with HGAP2. The resulting corrected reads were assembled into contigs with the Celera assembler. A final polishing step was then applied by aligning the raw reads back to the Celera contigs using BLASR and then passing them to Quiver to produce the final consensus sequence and derive PacBio quality scores.

Genome annotation and comparative genomic analysis were conducted with Rapid Annotation using Subsystems Technology (RAST)^123,124^. Annotations were made using the RASTtk annotation scheme with the Kmer dataset release 70 to annotate proteins and options to build a metabolic model. The *P. aeruginosa* M6A146 resistome was analyzed using the Resistance Gene Identifier (RGI) v5.2.1 via the web portal of the Comprehensive Antibiotic Resistance Database (CARD)^125^ v3.2.3.

### A 96-well microplate assay to identify spent media potentiating antibiotics

To facilitate medium-throughput assays using a 96-pin replicator, a set of 351 isolates (Supplementary Data 1) that grew in LB and BHI broths were arrayed into Nunc 96 Deepwell polystyrene plates (Thermo Fisher) containing 1.0 mL LB in each well, with arrangements as illustrated Fig. S12. These plates were incubated for 24 h at 37 °C and 250 RPM on an orbital shaker. Afterwards, 700 µl aliquots of these cultures were transferred using multichannel pipettes to another 96 Deepwell plate and combined with 300 µl of 50% w/v glycerol. These plates were sealed with VWR Aluminium Foil Seals for Cold Storage, and stored at −70 °C.

Spent media was produced in 96 Deepwell plates. Here, 1.0 mL aliquots of BHI were dispensed into 51 interior wells of 96 Deepwell plates with arrangements as illustrated in Fig. S12. Next, a 96-pin replicator was dipped in 95% ethanol and flamed to sterilize it. Subsequently, the 96-pin replicator was used to transfer small amounts of the frozen cultures stored at −70 °C in 96 Deepwell plates to the BHI in the freshly prepared plates. These plates were then sealed with Parafilm M Sealing Film and incubated for 24 h on an orbital shaker at 250 RPM and 25 or 37 °C, as dictated by the requirements the organisms (Supplementary Data 1). Following incubation, the 96 Deepwell plates were centrifuged at speeds from 3000 RPM up to 5,000 RPM for 10 to 20 min, as required, in a Sorvall LYNX 6000 Superspeed Centrifuge with a BioFLEX HC Swinging Bucket Rotor (ThermoFisher Scientific) to pellet the cells. Subsequently, 250 µl of spent media was transferred to a 96-well filtering plate with a 0.22 µm pore size and centrifuged at 3,000 × RPM for 10 min. Up to 750 µl of spent medium from each well was collected over three sequential centrifugal filtration steps.

The evaluation of spent media to potentiate meropenem (MEM) was predicated on a quantitative comparison between the growth of *P. aeruginosa* M6A146 in spent media blended with 50% v/v CA-MHB with MEM (potentiator assay) and without it (antibiosis assay) (Fig. 1). First, the potentiator assay was set up with 100 µl of CA-MHB + 4 µg/mL MEM (i.e., at twice the tested concentration) dispensed into the inner wells of a 96-well microplate (Fig. S12). A series of wells were reserved for growth and sterility controls (Fig. S12). Subsequently, 100 µl of filtered, spent medium from different test organisms was added to each of the wells (Fig. S12). Growth controls contained 100 µl spent medium from *P. aeruginosa* M6A146 that was combined with 50% v/v CA-MHB + 4 µg/mL MEM. Second, the antibiosis assay was set up in an identical manner except that the CA-MHB did not contain MEM. Potentiator and antibiosis assays were set up and executed in biological and technical triplicates.

To prepare an inoculum, *P. aeruginosa* M6A146 was streaked out twice in succession on LB + MEM 15 agar and each time grown overnight at 37 °C. Colonies from the second agar sub-culture were collected using a cotton swab and suspended in CA-MHB to match a McFarland 1.0 optical density standard. This suspension was then diluted thirtyfold in CA-MHB and verified by viable cell counting. A 5 µl aliquot of this standardized inoculum was added to each well of the potentiator and antibiosis assay plates, except for the sterility controls (Fig. S12). Microplates were sealed with Parafilm M Sealing film and incubated on an orbital shaker at 250 RPM and 37 °C for 24 h. Afterwards, the optical density at 600 nm (OD_600_) of these cultures was measured using an Enspire Multimode Plate Reader (Perkin-Elmer).

To evaluate the capacity of spent media to antagonize or potentiate the activity of meropenem against the target carbapenem-resistant *P. aeruginosa* strain, we performed a relative growth calculation. Here, we define relative growth (Eqn. 1) as the ratio of target strain growth (measured by OD_600_) in fresh broth mixed with the filtered, cell-free spent medium (SM) of another organism relative to its growth in that same mixture with meropenem (MEM). These calculations were normalized to the mean optical density (OD) for internal growth controls (GC) of the test strain and corrected for the mean background optical density of plastics and media in sterility controls (SC) within each microplate:

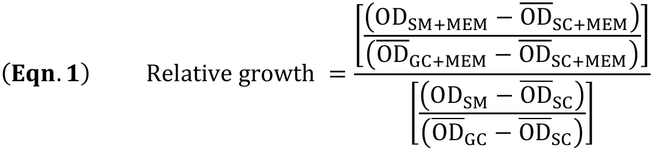

In simple terms, the numerator of this equation provides a measure for growth of the target strain in the presence of an antimicrobial in the spent medium of another organism, whereas the denominator provides a measure for growth of the target organism in the spent medium alone. Relative growth values of >1, ∼1, and <1 thus indicate antagonism, indifference, or synergism between the spent medium and target antimicrobial, respectively. Requirements for including data from individual microplates in the analysis were that sterility controls were negative for growth and that the OD_600_ of all growth controls were in the range of 0.2 to 0.65. Taken together, the relative growth calculation thus provides an unbiased way to identify organisms producing spent medium that alter the resistance of a target microbial strain to a test antimicrobial agent.

Secondary validation of hit organisms from the medium-throughput screen of spent media (Fig. 2) was carried out by evaluating meropenem potentiation by viable cell counts. Here, serial twofold dilution gradients of meropenem were prepared at twice the final tested concentration in CA-MHB. Subsequently, 100 µl of spent media, which was prepared as described above except for growing the hit organism in multiple wells of a 96 Deepwell plate, was added to each well of the microplate except for sterility controls. Microplates were sealed with Parafilm M Sealing Film and incubated for 22 h at 37 °C. Minimum inhibitory concentrations (MICs) were determined by measuring OD_600_ values for microplates in an Enspire Multimode Plate Reader, and killing was calculated from viable cell using established methods^112^.

### Additional susceptibility testing

Susceptibility of bacteria to purified, semi-synthetic or synthetic antibiotics and fatty amides was determined using standard methods for broth microdilution in 96-well microplates^126^ with minor modifications. Briefly, serial twofold dilution gradients of antibiotics were prepared at twice the final tested concentration in CA-MHB + 2% DMSO in 100 µl volumes. Subsequently, freshly prepared suspensions of CA-MHB + 2% DMSO with 320, 160, 80, or 40 µM fatty amides were dispensed in 100 µl volumes into wells, as required, to yield final in-plate concentrations of 160, 80, 40 and 20 µM these fatty amides, respectively. Growth and sterility controls contained 200 µl of CA-MHB + 2% DMSO only. In this fashion, the contents of every microplate well contained 2% DMSO. Recombinant human fatty acid amide hydrolase (FAAH, 2.5 units, Cayman Chemicals) was added to wells as indicated elsewhere in this report. Bacterial inoculum was prepared from strains that were streaked out twice on LB agar containing antibiotic selection, as required. Colonies from second agar sub-cultures were collected using a cotton swab and suspended in CA-MHB to match a McFarland 1.0 optical density standard. This suspension was diluted thirtyfold in CA-MHB, of which 5 µl were added to each well except for sterility controls. To minimize evaporation, the outermost wells of each plate were filled with 200 µl of sterile, ultrapure water. Microplates were sealed with Parafilm M Sealing Film and incubated for 22 h at 37 °C. MICs were determined by measuring OD_600_ values for microplates in an Enspire Multimode Plate Reader, and killing was calculated from viable cell counts as previously described^112^.

### Construction of allelic exchange vectors and deletion mutants

An in-frame deletion mutation in *ampC* was engineered in *P. aeruginosa* PAO1 using established protocols for two-step allelic exchange^63^. Briefly, primer pairs oRP_43/oRP_44 and oRP_45/oRP46 (Table S3) were used to amplify 488 bp and 564 bp DNA regions upstream and downstream of *ampC*, respectively. These PCR products were gel purified and then joined by splicing-by-overlap-extension (SOE) PCR using primers oRP_43 and oRP46, which had been tailed with *attB1* and *attB2* sequences, respectively. The SOE-PCR product was gel purified and then recombined with pDONRPEX18Gm using BP Clonase II. This reaction mixture was transformed into *E. coli* DH5α, plated onto LB + GEN 10 agar, and incubated overnight at 37 °C. Clones bearing the desired insert were identified via established protocols for colony PCR with M13F and M13R primers (oJJH367 and oJJH368, respectively, Table S3)^63,127^. The resulting allelic exchange vector, pENTRPEX18Gm::Δ*ampC* (pRP23, Table S2) was verified by Sanger sequencing using M13F and M13R primers. Subsequently, established methods for biparental mating^63^ with *E. coli* S17.1 (λ*_pir_*) transformed with pRP23 were used to introduce the vector into the recipient *P. aeruginosa* PAO1 strain (Table S1). Merodiploids were selected on VBMM + GEN 60 agar, and counter-selection was carried out on NSLB + 10% w/v sucrose agar, as previously described^63^. Sucrose-resistant, gentamicin-sensitive *P. aeruginosa* strains were verified by replicate plating on LB agar, LB + GEN 60 agar, and NSLB + 10% w/v sucrose agar. The strain with the desired deletion mutation was identified via colony PCR with the primers oRP_43 and oRP46, and the PCR product was verified by Sanger sequencing with those same primers, yielding *P. aeruginosa* RP146 (Table S1).

### Construction of miniTn7 vectors and transposon mutagenesis

A *P. aeruginosa* strain was designed and built to enable inducible expression of *ampC*. To begin, DNA encoding *araC-P_BAD_::ampC* was assembled and fused with the backbone of pUC18miniTn7T2.1-Gm-GW (Table S2). Here, the *ampC* open reading frame (ORF) was cloned from *P. aeruginosa* PAO1 gDNA with primers oRP_51 and oRP_52 (Table S3). The oRP1_51 primer was tailed with an artificial ribosomal binding site, 5’-GAGGAGGATATTC-3’, as previously described^128^. In parallel, a DNA fragment encoding *araC-P_BAD_* was cloned from pJN105^129^ (Table S2) using the primers oH229 and oH230 as previously described^111^, while the backbone of pUC18miniTn7T2.1-Gm-GW^130^ was cloned with primers oRP92 and oRP93 (Table S3). All primers were tailed with homology extensions (Table S3), which were used to facilitate Gibson assembly^131^ of the 3 PCR products, which had been gel purified. Subsequently, Gibson reaction mixtures were transformed into *E. coli* DH5α by electroporation, which was then spread onto LB + GEN 10 agar and incubated overnight at 37 °C. Colony PCR with primers oRP_51 and oRP_52 was used to identify clones bearing the desired insert, and the resulting plasmid, pRP1 (Table S2), was verified via Sanger sequencing using primers oRP_51 and oRP_52. Mutagenesis of *P. aeruginosa* RP146 with pRP1 was executed via established protocols^64^ for miniTn7 mutagenesis with the helper plasmid pTNS3 (Table S2). Transposon mutants were selected on LB + GEN 30 agar. The resulting strain, *P. aeruginosa* PAO1 Δ*ampC attTn7*::miniTn7T2.1-*araC-P_BAD_::ampC* (RP20, Table S1), was thus engineered for L-(+)-arabinose-inducible expression of *ampC*.

### Ethidium bromide accumulation assay

Bacterial strains were streaked out twice on LB agar, and an inoculum was prepared by suspending colonies from the second agar subculture in PBS to attain an OD_600_ = 0.3. Assays were set up in clear-bottomed, black-walled microplates. In technical triplicate, 4.1 μl of 1 mM oleamide, 1 mM elaidamide or DMSO was added to 195.9 μl of the inoculum. By contrast, 4 μl of 50 µg/mL PAβN was added to 196 μL of inoculum, or 0.1 μl of 0.5 μg mL^-1^ CST, was added to 199.9 μl of inoculum. Finally, 2 μl of 1 µg/mL ethidium bromide was dispensed into every well. Fluorescence measurements using excitation and emission wavelengths of 532 and 572 nm, respectively, were recorded every 5 min for 60 min using an Enspire Multimode Plate Reader (Perkin-Elmer).

### Subcellular fractionation

Periplasmic fractions of *P. aeruginosa* were obtained using established protocols for differential centrifugation^132^. Here, *P. aeruginosa* was grown in 500 mL of LB in a baffled 1 l flask for 16 - 18 h at 37 °C and 250 RPM. Bacterial cells were collected by centrifugation at 5,000 × g for 20 min at 4 °C. The supernatant (e.g., spent media) was removed and retained for further testing. The cell pellet was suspended in 5 mL of buffer (containing 0.2 M Tris-HCl, 1 mM EDTA, 1 M sucrose, adjusted to pH 8.0) containing 12 mg/mL lysozyme (ThermoFisher Scientific). This mixture was incubated for 10 min at room temperature. Subsequently, 20 mL ultrapure water was added to the reaction, and it was placed on ice for 20 min. This mixture was centrifuged at 100,000 × g for 2 h at 4 °C, yielding a supernatant containing the periplasmic fraction of cells, which was kept for analysis. Protein concentrations in periplasmic fractions were determined using the Pierce 660 nm assay.

### Nitrocefin assays

A 1 mM nitrocefin stock solution was prepared by dissolving 1 mg nitrocefin (Sigma-Aldrich) in 100 µl DMSO and adding 1.9 mL PBS to it. Assays for β-lactamase activity were set up in clear-bottomed, black-walled microplates. To assess any direct effects of oleamide on β-lactamase activity, assays were set up with 175.9 µl of a periplasmic fraction prepared from *P. aeruginosa* cells as described above, 20 µl of the 1 mM nitrocefin stock solution, and either 4.1 µl of DMSO or 4.1 µl of 0.98 mM oleamide in DMSO. Absorbance at 490 nm (A_490_) was measured every minute for 60 min using an Enspire Multimode Plate Reader (Perkin-Elmer).

### Light microscopy and image analysis

*P. aeruginosa* M6A146 was grown in microplate wells using the conditions above for additional susceptibility testing. A total of 2 mL of bacteria from each treatment group (DMSO, + 20 µM oleamide, or + 20 µM elaidamide) were pooled. Cells were collected by centrifugation at 9400 × g for 2 min, washed twice with 1 mL PBS, and suspended in 800 μl of PBS. Next, 2.5 μl of a Syto 9 and propidium iodide (PI) solutions (from the LIVE/DEAD BacLight Bacterial Viability Kit, ThermoFisher Scientific) were added to the cells, which were incubated in the dark for 15 min. Afterwards, 5 μl of the dye-stained sample was transferred onto a 1 cm^2^ agar pad in a microscope chamber as per established techniques^133^. Cells were imaged using a Nikon A1R confocal microscope using a 20×/0.7 numerical aperture objective. Samples were excited with the 488 nm laser, and emission was collected using the green (550 – 500 nm) and red (570 – 620 nm) channels, corresponding to Syto 9 and PI, respectively.

ImageJ (v1.53t) was used to open the Nikon (.nd2) files, and the integrated auto-threshold (triangle) function was used to select pixels in the Syto 9 and PI channels that exceeded the background^134^. We then applied a composite mask (Boolean AND) to select all pixels with non-zero intensities in both the Syto 9 and PI channels. Subsequently, the intensity values from each channel were then used to calculate a ratiometric Syto 9/PI score for each pixel in the image, using the following equation:

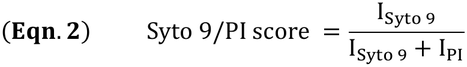

Here, I_Syto 9_ is the intensity of the pixel in the Syto 9 channel and I_PI_ is the intensity of the pixel in the PI channel. The score ranges between 0 and 1 for each pixel; a higher score indicates more Syto 9 signal, a lower score indicates more PI signal; a score of 0.5 indicates equal amounts of Syto 9 and PI signals.

### Standard chemical methods

All solvents for the isolation and synthesis of N-acyl amides were ACS grade. ^1^H NMR spectra were recorded on a Bruker Avance III 400 instrument at 400 MHz and a Bruker Avance 400 instrument at 400 MHz using standard 1D pulse programs for samples dissolved in CD_3_OD or CDCl_3_ (Sigma Aldrich), where all NMR spectra were referenced to the residual protonated solvent signal (δ_H_ 3.31 and 7.26 ppm, respectively). High resolution electrospray ionization mass spectrometry (HR-ESI-MS) was performed on an Agilent Q-TOF 6520 mass spectrometer with gas chromatography-mass spectrometry (GC-MS) performed on an Agilent HP5975 instrument. Normal phase flash column chromatography was performed on a Biotage Isolera One Dalton using 12g Silicycle HP silica columns (column volume of 15 mL) at a flow rate of 12 mL min^-1^ with UV monitoring from 200 to 800 nm. Size exclusion chromatography (SEC) was performed on a GE AKTA Pure 25 M1 system using a Supered 30 Increase 10/300 column (column volume of 23.562 mL) at a flowrate of 0.8 mL min^-1^ with UV monitoring at 215, 254 and 280 nm. Semi-preparative reversed phase high performance liquid chromatography (RP-HPLC) was performed using an Agilent Infinity 1260 HPLC system, with a Phenomenex Kinetex EVO C18 silica column (5 μm, 100 Å, 250 × 10 mm), equipped with a Phenomenex Kinetex Core-Shell EVO C18 silica guard column (sub-2 μm, 10 × 10 mm ID) and an Agilent 1260 G1365C multiple wavelength detector (220 nm, 254 nm, 280nm, & 365 nm) at a flow rate of mL min^-1^. Normal phase thin layer chromatography (TLC) was performed on aluminum backed TLC plates (200 μm thickness, Millipore). TLC plates were visualized under UV (254 nm) and by dipping plates in a solution of phosphomolybdic acid followed by charring.

### Bioactivity-guided fractionation

Lyophilized chemical extracts and fractions were dissolved in DMSO to provide a standardized 5 mg/mL concentration. Each sample was tested at a final concentration of 100 µg/mL. Microplate susceptibility testing of *P. aeruginosa* M6A146 to twofold dilutions of meropenem in combination with these extracts was otherwise executed as described above for susceptibility testing. Changes in antibiotic sensitivity were ascertained by comparing OD_600_ values for wells containing DMSO alone to those wells additionally containing the reconstituted, purified chemical fraction.

### Isolation of oleamide

Spent media of *N. perflava* RP9 (5 L) was first extracted with 3 × 300 mL hexanes, followed by equivalent washes of CH_2_Cl_2_, EtOAc and n-BuOH. The hexanes extract (16 mg) was found to possess adjuvant activity with meropenem and was subsequently fractionated by normal phase flash column chromatography using a stepwise gradient from 100% hexanes to 1:1 hexanes/EtOAc in 10% increments (5 CV/step), followed by 100% EtOAc (5 CV), 1:1 EtOAc/MeOH (5 CV) and 100% MeOH (5 CV) washes to give oleamide (3.1 mg). All spectroscopic and spectrometric data were consistent with that of a commercial standard.

Spent broth of *Aeromonas* sp. CCT22 (1 L) was first extracted with 3 × 300 mL hexanes, followed by equivalent washes of CH_2_Cl_2_, EtOAc and *n*-BuOH to give oleamide (4.5 mg) from the hexanes extract. All spectroscopic and spectrometric data were consistent with that of a commercial standard.

Spent broth of *S. lugdunensis* 47 (1 L) was extracted using HP-20 Diaion^®^. The HP-20 (200.03 g) was initially activated with MeOH (2 × 500 mL) for 30 min then rinsed with H_2_O (2 L) by vacuum filtration. The spent broth was then added to the activated HP-20 and stirred together at room temperature for 2h, whereby the broth was then removed from the resin by vacuum filtration and concentrated *in vacuo* to give a H_2_O extract (3.16 g). MeOH (2 × 500 mL) was added to the HP-20 and stirred together for 30 min at room temperature, then removed from the resin by vacuum filtration and concentrated *in vacuo* to give a MeOH extract (11.05 g). Isopropanol (2 × 500 mL) was then added to the HP-20 and stirred together for 30 min at room temperature, then removed from the resin by vacuum filtration and concentrated *in vacuo* to give an isopropanol extract (214 mg). The isopropanol extract showed adjuvant activity with a portion of the extract (20.0 mg) fractionated by size exclusion chromatography using 7:3 CH_3_CN/H_2_O to give seven fractions. Fraction 2 was subsequently fractionated by RP-HPLC using a linear gradient (0-5 min – 95:5 H_2_O/CH_3_CN buffered with 0.1% TFA; 5-25 min – linear gradient from 95:5 to 55:45 H_2_O/MeOH) to give 6 fractions. Fraction 6 was identified as oleamide (1.0 mg), with all spectroscopic and spectrometric data being consistent with that of a commercial standard.

### Acquisition and/or synthesis of fatty acids and fatty amides

Fatty acids were purchased from Combi-Blocks and Sigma Aldrich as required. A commercial standard of oleamide was purchased from Combi-Blocks Incorporated. Primary fatty amides were synthesized in-house from fatty acids as summarized below:

#### Stearamide

Stearic acid (0.5 g) was refluxed with thionyl chloride (approximately 3 mL) for 2 h. Excess thionyl chloride was removed by rotary evaporation. The resulting acyl chloride was cooled in an ice bath with subsequent dropwise addition of concentrated aqueous ammonia (7.5 mL). The resulting white powder was vacuum filtered and dried under reduced pressure. The precipitate was dissolved in CHCl_3_ and purified by normal phase column chromatography (silica gel column using a 5% methanol: chloroform mobile phase). The resulting crystals were re-concentrated and dissolved in a 1:1 mixture of ether:CH_2_Cl_2_ and partitioned with NaOH (1M, approximately 2 mL). The CH_2_Cl_2_ layer was concentrated to dryness yielding a white solid. Comparison of spectroscopic and spectrometric data with literature values confirmed the production of stearamide (35 mg, 7%).

#### Palmitamide

Palmitic acid (0.5 g) was refluxed with thionyl chloride (approximately 3 mL) for 2h. Excess thionyl chloride was removed by rotary evaporation. The resulting acyl chloride was cooled in an ice bath with subsequent dropwise addition of 7.5 mL of concentrated aqueous ammonia (7.5 mL). The white precipitate was vacuum filtered and dried under reduced pressure. The precipitate was dissolved in CHCl_3_ and CH_2_Cl_2_ before addition of H_2_O, 1M NaOH, and a saturated NaCl solution. The organic layer was extracted and dried. The resulting white precipitate was purified by recrystallization with chloroform. Comparison of spectroscopic and spectrometric data with literature values confirmed the production of palmitamide (168 mg, 34%).

#### Arachidiamide

Arachidic acid (0.5 g) was refluxed with thionyl chloride (approximately 3 mL) for 2h. Excess thionyl chloride was removed by rotary evaporation. Cold concentrated aqueous ammonia was added dropwise to the flask yielding a white precipitate. The precipitate was vacuum filtered and dried under reduced pressure. Inorganic salts were removed by hot filtration via a syringe. The remaining liquor was concentrated and extracted with CH_2_Cl_2_, H_2_O, 1M NaOH, and a saturated NaCl solution. The final product was purified by recrystallization with CHCl_3_. Comparison of spectroscopic and spectrometric data with literature values confirmed the production of arachidiamide (30 mg, 6%).

#### Elaidamide

Elaidic acid (0.5 g) was dissolved in minimal CH_2_Cl_2_ and DMF (2 drops), cooled on ice, with oxalyl chloride (3 equivalents) subsequently added. The mixture was brought to room temperature prior to reflux for 2h. Excess solvent and oxalyl chloride was removed by rotary evaporation. The product was cooled on ice with concentrated aqueous ammonia (7 mL) was added dropwise. The white solid was vacuum filtered and dried under reduced pressure, redissolved in CH_2_Cl_2_ and extracted with H_2_O, 1M NaOH, and a saturated NaCl solution. The organic layer was concentrated to dryness. Comparison of spectroscopic and spectrometric data with literature values confirmed the production of elaidamide (220 mg, 44%).

#### Arachidonamide

Arachidonic acid (151.0 mg) was dissolved in minimal CH_2_Cl_2_ and DMF (2 drops), cooled on ice, with oxalyl chloride (0.1 mL) subsequently added. The mixture was brought to room temperature prior to reflux for 2h. Excess solvent and oxalyl chloride was removed by rotary evaporation. The product was cooled on ice with concentrated aqueous ammonia (5 mL) was added dropwise. The brown oil was partitioned with CH_2_Cl_2_, ammonia, H_2_O, and a saturated NaCl solution. The crude product was purified by column chromatography (EtOAc) and liquid-liquid partition (CH_2_Cl_2_/H_2_O) and was concentrated to dryness under reduced pressure. Comparison of spectroscopic and spectrometric data with literature values confirmed the production of arachidonamide (50 mg, 33%).

#### Linoleamide

Linoleic acid (530.0 mg) was dissolved in minimal CH_2_Cl_2_ and DMF (2 drops), cooled on ice, with oxalyl chloride (approximately 0.5 mL) was subsequently added dropwise. The mixture was then refluxed on low heat for 2h. Excess solvent and oxalyl chloride was removed by rotary evaporation. The product was cooled on ice with concentrated aqueous ammonia (5 mL) was added dropwise. The solidified oil was dissolved in CH_2_Cl_2_ and partitioned with H_2_O, NaCl, and 0.1M NaOH with the organic layer purified by normal phase column chromatography (EtOAc containing 0.1% Et_3_N). Fractions containing the amide were concentrated and dried under reduced pressure. Comparison of spectroscopic and spectrometric data with literature values confirmed the production of linoleamide (143 mg, 27%).

#### Palmitolamide

Palmitoleic acid (100.0 mg) was dissolved in minimal CH_2_Cl_2_ and DMF (2 drops), cooled on ice, with oxalyl chloride (approximately 0.1 mL) was subsequently added dropwise. The mixture was then refluxed for 2h. Excess solvent and oxalyl chloride was removed by rotary evaporation. The product was cooled on ice with concentrated aqueous ammonia (approximately 2 mL) was added slowly. The solidified oil was vacuum filtered and washed with excess ammonia. The solidified oil was partitioned between CH_2_Cl_2_ and a 0.1M NaOH/saturated NaCl solution. The organic layer was dried over Na_2_SO_4_ and concentrated. Comparison of spectroscopic and spectrometric data with literature values confirmed the production of palmitolamide (57 mg, 57%).

#### Oleamide

Oleic acid (522.0 mg) was dissolved in minimal CH_2_Cl_2_ and DMF (2 drops), cooled on ice, with oxalyl chloride (approximately 0.45 mL) was subsequently added dropwise. The mixture was then refluxed for 2h. Excess solvent and oxalyl chloride was removed by rotary evaporation. The product was cooled on ice with concentrated aqueous ammonia (approximately 2 mL) was added slowly. The yellow solidified oil-like material was vacuum filtered and washed with excess ammonia. The filtered material was dissolved in CH_2_Cl_2_ and partitioned with a 0.1M NaOH/saturated NaCl solution. The organic layer was dried over Na_2_SO_4_ and concentrated. The product was purified by normal phase column chromatography (EtOAc containing 0.1% Et_3_N). The resulting fractions were combined and concentrated, where comparison of spectroscopic and spectrometric data with literature values confirmed the production of oleamide (200 mg, 38%).

#### Linolenamide

Linolenic acid (100.0 mg) was dissolved in minimal CH_2_Cl_2_ and DMF (2 drops), cooled on ice, with oxalyl chloride (approximately 0.1 mL) was subsequently added dropwise. The mixture was then refluxed for 2h. Excess solvent and oxalyl chloride was removed by rotary evaporation. The product was cooled on ice with concentrated aqueous ammonia (approximately 2 mL) was added slowly. The solidified orange-yellow material was vacuum filtered and washed with excess ammonia. The filtered material was dissolved in CH_2_Cl_2_ and partitioned with a 0.1M NaOH/saturated NaCl solution. The organic layer was dried over Na_2_SO_4_ and concentrated. The product was purified by normal phase column chromatography (7% MeOH in EtOAc). The resulting fractions were combined and concentrated, where comparison of spectroscopic and spectrometric data with literature values confirmed the production of linolenamide (49 mg, 49%).

#### cis-Vaccenamide

cis-Vaccenic acid (50.0 mg) was dissolved in minimal CH_2_Cl_2_ and DMF (2 drops), cooled on ice, with oxalyl chloride (0.05 mL) was subsequently added dropwise. The mixture was then refluxed for 2h. Excess solvent and oxalyl chloride was removed by rotary evaporation. The product was cooled on ice with concentrated aqueous ammonia (approximately 2 mL) was added slowly. The white solidified oil-like product was vacuum filtered and washed with excess ammonia. The filtered material was dissolved in CH_2_Cl_2_ and partitioned with a 0.1M NaOH/saturated NaCl solution. The organic layer was dried over Na_2_SO_4_ and concentrated. The resulting fractions were combined and concentrated, where comparison of spectroscopic and spectrometric data with literature values confirmed the production of cis-vaccenamide (20 mg, 40%).

### Preparation of liposomes mimicking the *P. aeruginosa* inner membrane

Lyophilised 1-palmitoyl-2-oleoyl-*sn*-glycero-3-phosphoethanolamine (POPE) and 1-palmitoyl-2-oleoyl-*sn*-glycero-3-phospho-(1′-rac-glycerol) (POPG) were purchased from Avanti Polar lipids (Alabaster, Al). These lipid powders were used without further purification. The fluorescent dye 6-dodecanoyl-2-dimethylaminonaphthalene (laurdan GP) was purchased from Molecular Probes. Stock solutions were prepared by dissolving this dye in ACS grade chloroform (Sigma Aldrich).

A model membrane mimic for *P. aeruginosa* was made by preparing a binary mixture of POPE:POPG at a 4:1 molar ratio. POPE and POPG are highly abundant in the cell membrane of *P. aeruginosa*. Lipid stock solutions were made by dissolving the lipids in a 7:3 chloroform:methanol mixture with laurdan GP in a 1:500 dye-to-lipid molar ratio. Lipid films were re-hydrated with buffer containing 20 mM 4-(2-hydroxyethyl)-1-piperazineethanesulfonic acid (HEPES, Sigma-Aldrich) and 100 mM NaCl (pH 7.4). Vortex mixing, sonication, and at least 5 freeze-thaw cycles were then used to generate multilamellar vesicles. Large unilamellar vesicles were prepared by passing the multilamellar vessicles through a Whatman Nuclepore polycarbonate filter (100 nm pore size, Sigma-Aldrich) using a Mini Extruder (Avanti). The multilamellar suspension was passed between 2 gas tight syringes ≥20 times to ensure uniform size distribution of large unilamellar vesicles. For each lipid system, multilamellar vessicles were extruded above the phase transition temperature (T_m_). After extrusion, phospholipid concentration was determined using a phosphate assay according to Ames.

### Generalized polarization test

Fluorescence measurements for laurdan GP were performed using a Cary Eclipse fluorimeter (Agilent technologies, Santa Clara, CA) at an excitation wavelength of 340 nm. Emission wavelengths were measured at both 440 and 490 nm as an average of 3 independent measurements. We used an excitation and emission band pass of 5 nm. Samples consisted of 0.3 mM large unilamellar vesicles in small volume quartz cuvettes (Starna Scientific) with 0.15 – 1.5 mM of oleamide or elaidamide. A negative control with DMSO was tested to ensure that the solvent used to dissolve the fatty amides did not significantly impact model membrane properties. Fatty amides were added to the large unilamellar vesicles and allowed to incubate for 5 minutes at room temperature prior to any measurements. Temperature was controlled to ± 0.1 °C using a circulating water bath (Agilent Technologies).

### Mouse wound infection model

SKH1-elite outbred hairless mice (strain code 477) were purchased from Charles River Laboratories and bred in a specific pathogen-free facility. An excision wound model was modified from a previous study by our group^52^. Briefly, 10 – 12 week-old male and female mice were selected randomly for treatment groups. General anesthesia was induced using 4% isoflurane at a flow rate of 1 L/min in a 95% O_2_ – 5% CO_2_ mixture; 1 – 3% isoflurane was used to maintain anesthesia. A 4 mm full thickness circular excision wound was created on the left flank and infected with ∼5 × 10^7^ CFU of *P. aeruginosa* MB6476 and covered with an occlusive dressing. Mice were administered 5 mg/kg meloxicam once daily for 3 days via subcutaneous injection for pain relief. Mice were treated topically with the vehicle (100% DMSO), 3% oleamide and/or 8 µg/ml imipenem, twice daily commencing 1 h post-infection. On day 4 post-infection, mice were euthanized, and the wound tissue, kidneys and spleen were harvested and homogenized for serial dilutions and bacterial viable cell counts.

## RESOURCE AVAILABILITY

### Lead contact

- Requests for further information and resources should be directed to and will be fulfilled by the lead contact, Dr. Joe Harrison.

### Materials availability

- All microbial isolates are cataloged in and are available from the Alberta Microbiota Repository (AMBR) at the University of Calgary.
- All plasmids and genetically modified bacterial strains are available from the lead contact without restriction.
- All other unique/stable reagents generated in this study are available from the lead contact without restriction.

## Data and code availability

- The complete *P. aeruginosa* M6A146 genome sequence was deposited in National Center for Biotechnology Information (NCBI) Genbank with accession CP113974.1 and appears in the *Pseudomonas* genome database with accession GCF_026870135.1. Sequencing data were deposited with links to BioProject accession number PRJNA909305 in the NCBI BioProject database.
- The data specification for capturing strain information in DataHarmonizer is available on Github (https://github.com/cidgoh/AMBR_Contextual_data_Specification).
- All data reported in this paper will be shared by the lead contact upon request.
- Any additional information required to reanalyze the data reported in this paper is available from the lead contact upon request.

## ACKNOWLEDGMENTS

JJH has been supported by a Tier II Canada Research Chair from the Canadian Institutes of Health Research (CIHR), a Discovery Grant from the Natural Sciences and Engineering Research Council (NSERC) of Canada, and the Leaders’ Opportunity Fund of the Canada Foundation for Innovation (CFI). DJD and JJH have received seed funding from the Experimental Therapeutics program at the Arnie Charbonneau Cancer Institute. JMC, MDP, DJM, DJD and JJH have received strategic funding from the Vice President of Research at the University of Calgary. KDM and JJH received funding from the Antimicrobial Resistance – One Health Consortium, which is supported by the Major Innovation Fund program of the Ministry of Jobs, Economy, and Innovation (JEI), Government of Alberta. Lastly, KDM and JJH received funding from the CIHR Canadian Microbiome Initiative 2: Research Core program, which supports Integrated Microbiome Platforms for Advancing Causation Testing and Translation (IMPACTT). The authors thank Rayna Patel, Tie Wang, and Chloe Lee for technical assistance as well as the Live Cell Imaging Laboratory at the University of Calgary.

## AUTHOR CONTRIBUTIONS

Conceptualization: JMC, DJM, DJD and JJH. Methodology: RP, AJF, DJM, DJD, JJH. Software: EJG, NSJ, CB, RC, DD, WH. Validation: YL, SP, KDM, AK. Formal analysis: RP, AJF, CB, PC, EJP, SER, JJH. Investigation: RP, AJF, SA, NBT, BR, KC, AJG, YL, SP, CB, ZT, JDL, MA-S. Resources: RP, SA, NKM, VA, CT, BW, MDP, AK, PC, EJP, WH, KDM, DJM, DJD, JJH. Data curation: RP, SA, VA, EJG, WH, JJH. Writing – Original Draft: RP and JJH. Writing – Review and Editing: RP, VA, EJG, JMC, AK, PC, EJP, WH, KDM, SER, DJD, JJH. Visualization: RP and JJH. Supervision: MDP, AK, PC, EJP, WH, KDM, DJM, SER, DJD, JJH. Project administration: JJH. Funding acquisition: WH, KDM, DJM, DJD, JJH.

## DECLARATION OF GENERATIVE AI AND AI-ASSISTED TECHNOLOGIES

The authors have not used AI in the preparation of this work.

## SUPPLEMENTAL INFORMATION

**Document S1. Figures S1–S12, and Tables S1–S5**

**Data S1.** Strains isolated by massively parallel culturing in this study and their contextual metadata.

**Data S2.** Mean relative growth measurements for *P. aeruginosa* M6A146 in spent media with and without the addition of meropenem.

