## Supplemental information for "Antibiotic potentiators from microbiota that permeate the *Pseudomonas aeruginosa* cell envelope"

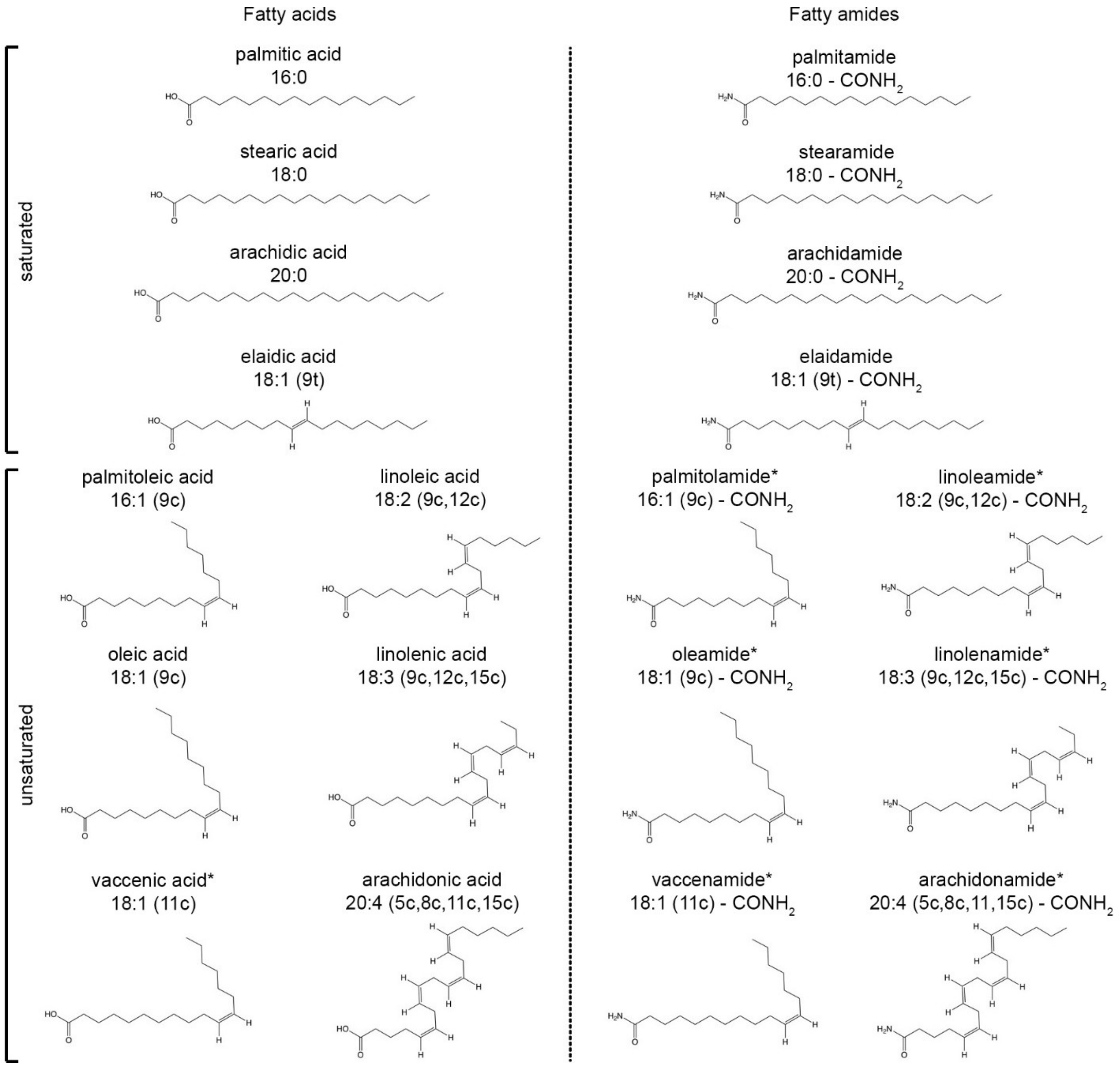


**Fig. S1.** Structures, naming and notation of the fatty acids and fatty amides used in this study. An asterisk denotes a molecule that increases the susceptibility of carbapenem-resistant *P. aeruginosa* M6A146 to the last resort antibiotic, meropenem.

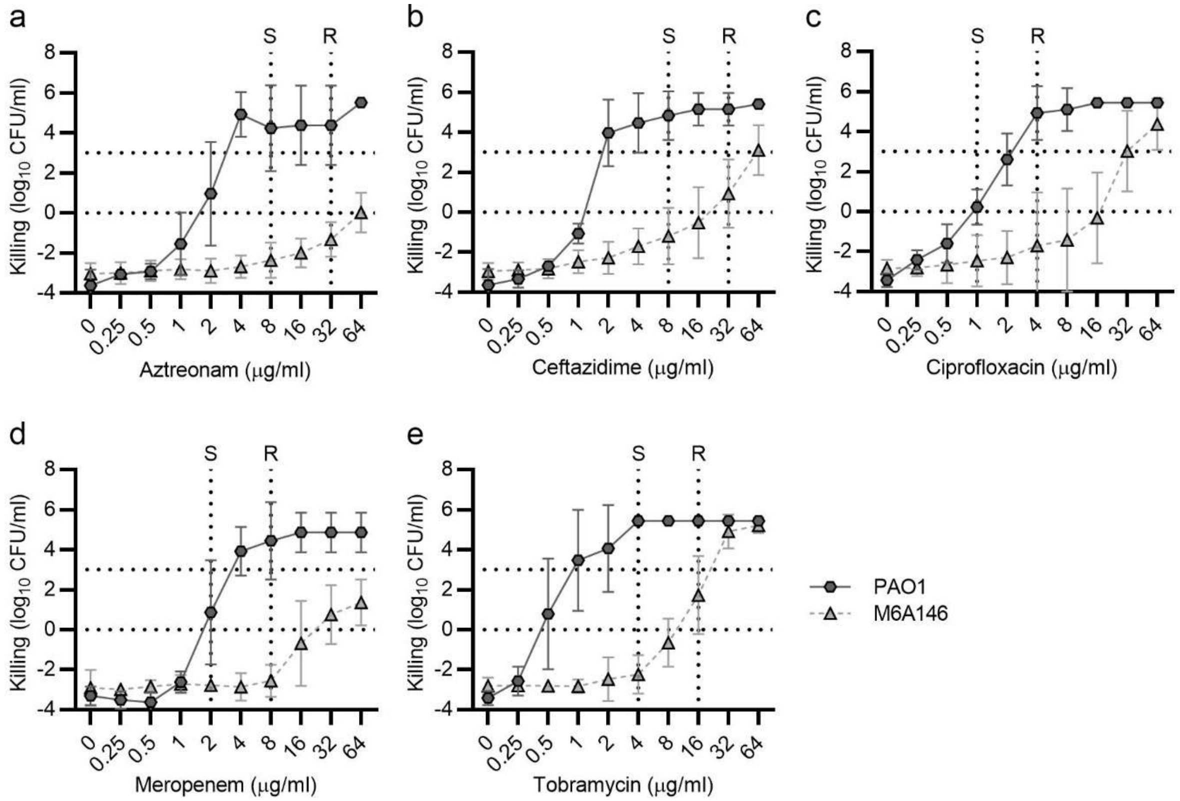


**Fig. S2.** *Pseudomonas aeruginosa* M6A146 is a multidrug-resistant clinical isolate. Susceptibility of *P. aeruginosa* M6A146 to (a) aztreonam, (b) ceftazidime, (c) ciprofloxacin, (d) meropenem, and (e) tobramycin. Points and error bars represent the means and standard deviations of 9 replicates (3 technical replicates from 3 independent biological replicates). Clinical breakpoints for sensitivity (S) and resistance (R) [ref. [1]] are denoted with vertical dotted lines. Horizontal dotted lines at 0 and 3 $log_{10}$ CFU/mL denote inhibitory and bactericidal thresholds, respectively.

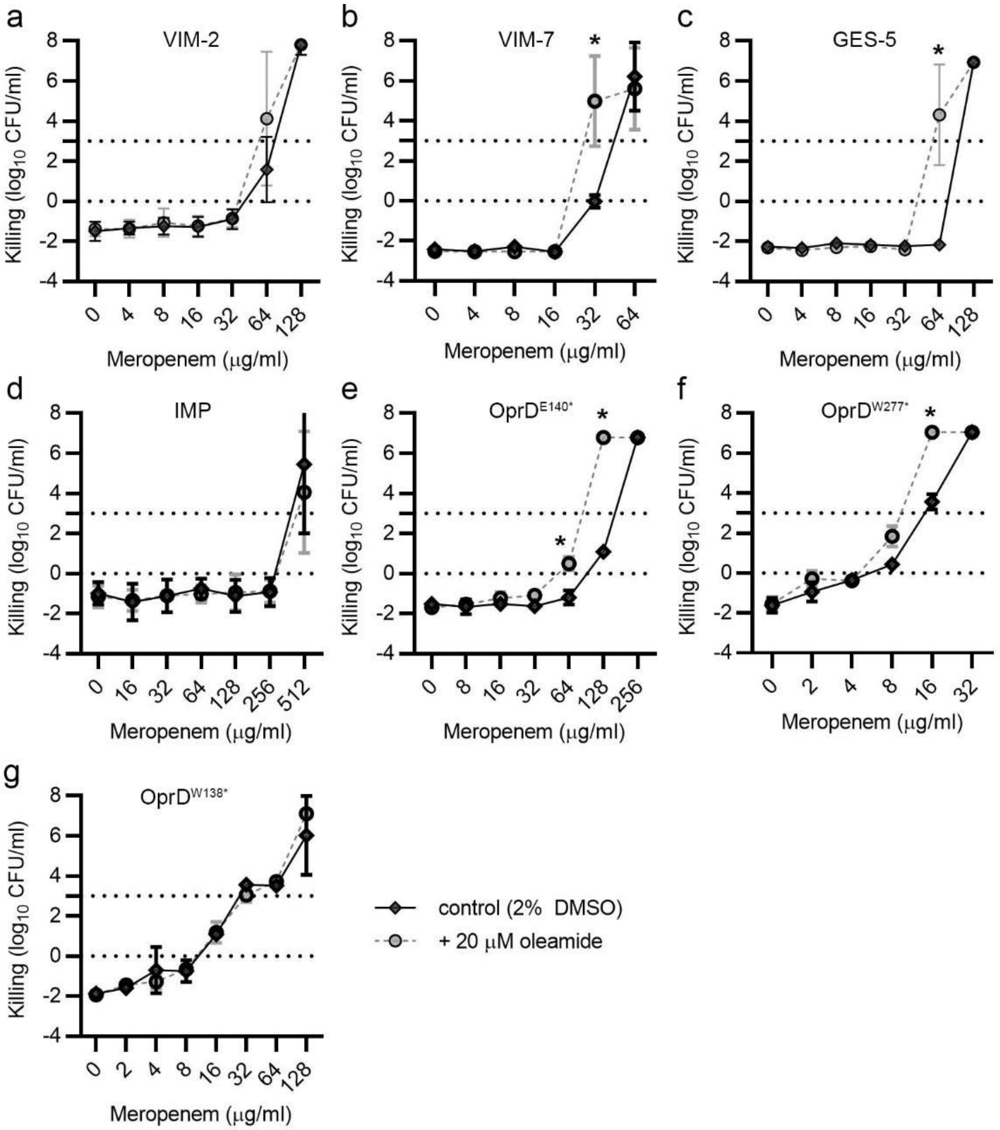


**Fig. S3.** Oleamide potentiates meropenem against many *P. aeruginosa* isolates expressing different carbapenem resistance determinants. Strain details appear in Table S1. Meropenem susceptibility of (a) *P. aeruginosa* AFC-7 Kenya expressing the Verona integron–encoded metallo-β-lactamase (VIM) 2, (b) *P. aeruginosa* JJH1352 expressing VIM-7, (c) *P. aeruginosa* JJH1353 expressing Guiana extended spectrum 5 (GES-5) β-lactamase, (d) *P. aeruginosa* JJH1351 expressing imipenemase (IMP), (e) *P. aeruginosa* JJH1355 expressing $OprD^{W65*}$, (f) *P. aeruginosa* JJH1357 expressing $OprD^{W277*}$, and (g) *P. aeruginosa* JJH1355 expressing $OprD^{W138*}$. Points and error bars represent the means and standard deviations of ≥3 independent biological replicates. Significant differences were calculated using a two-tailed Student's t-test with the Holm-Sidak multiple test correction (* denotes $p < 0.05$). Horizontal dotted lines at 0 and 3 $log_{10}$ CFU/mL denote inhibitory and bactericidal thresholds, respectively.

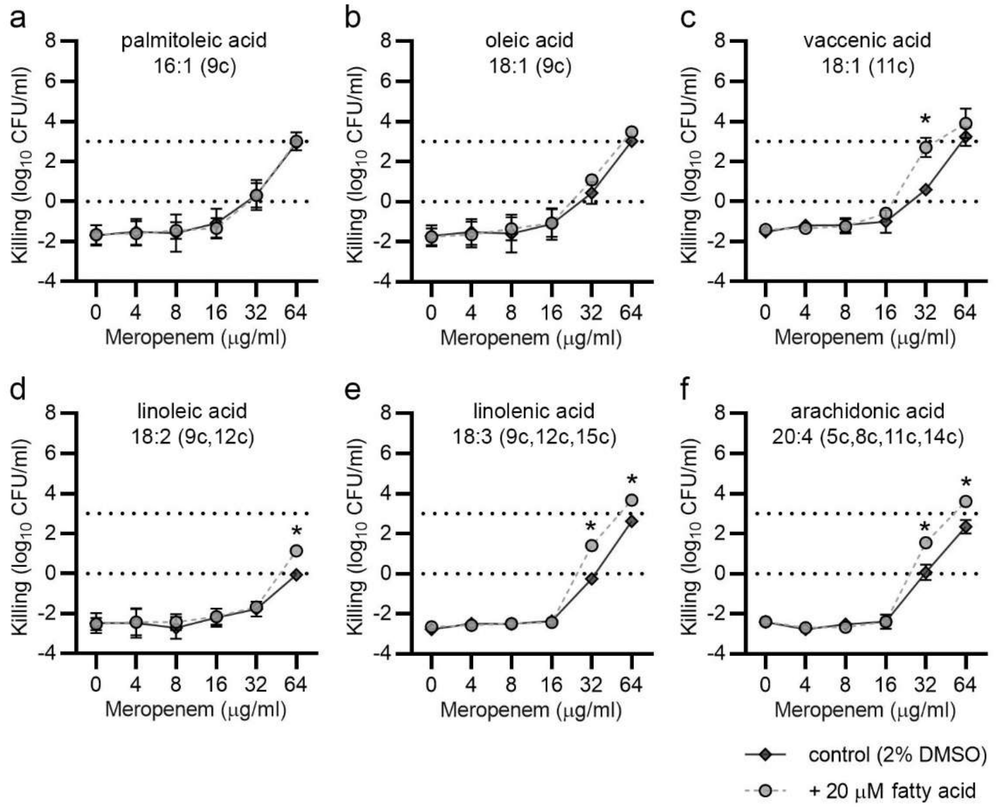


**Fig. S4.** Some unsaturated fatty acids have a significant but small effect on the susceptibility of carbapenem-resistant *P. aeruginosa* M6A146 to meropenem. Meropenem susceptibility of *P. aeruginosa* M6A146 in (b) palmitoleic acid, (b) oleic acid, (c) vaccenic acid, d) linoleic acid, (e) linolenic acid, and (f) arachidonic acid. Points and error bars represent the means and standard deviations of ≥3 independent biological replicates. Significant differences were calculated using a two-tailed Student's t-test with the Holm-Sidak multiple test correction (* denotes $p$ < 0.05). Horizontal dotted lines at 0 and 3 $\log_{10}$ CFU/mL denote inhibitory and bactericidal thresholds, respectively.

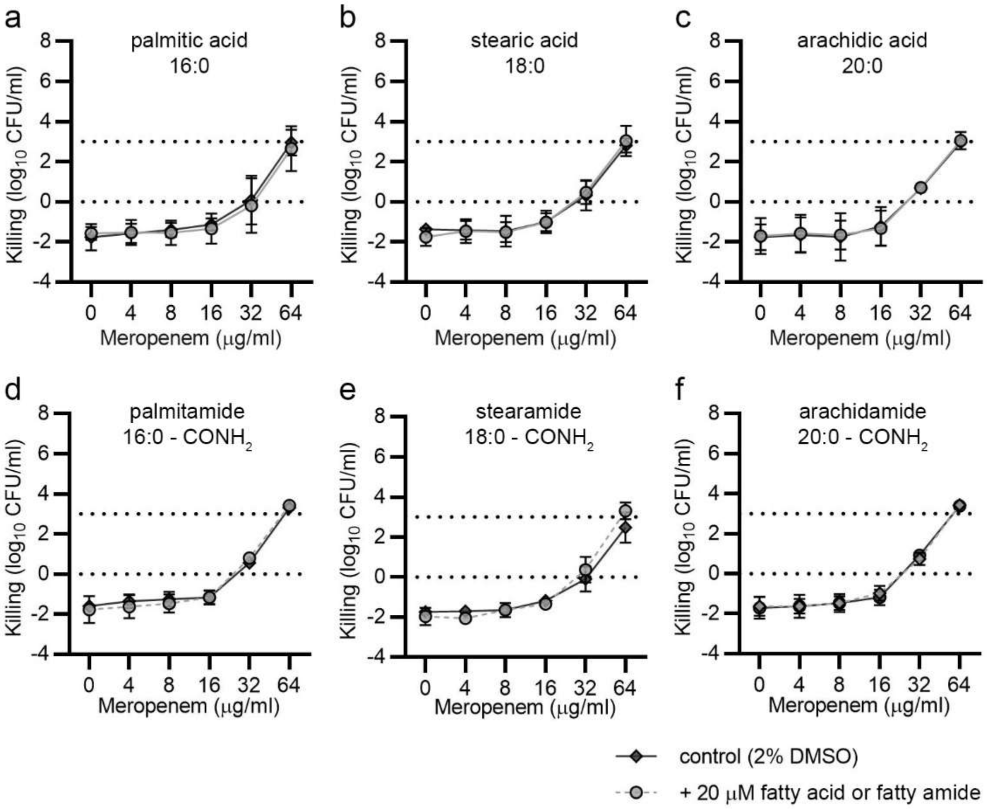


**Fig. S5.** Saturated fatty acids and saturated fatty amides do not affect the susceptibility of carbapenem-resistant *P. aeruginosa* M6A146 to meropenem. Meropenem susceptibility of *P. aeruginosa* M6A146 in (b) palmitic acid, (b) stearic acid, (c) arachidic acid, d) palmitamide, (e) stearamide, and (f) arachidamide. Points and error bars represent the means and standard deviations of ≥3 independent biological replicates. There were no significant differences when controls and fatty acid- or fatty amide-treated groups were compared using a two-tailed Student's t-test with the Holm-Sidak multiple test correction. Horizontal dotted lines at 0 and 3 $log_{10}$ CFU/mL denote inhibitory and bactericidal thresholds, respectively.

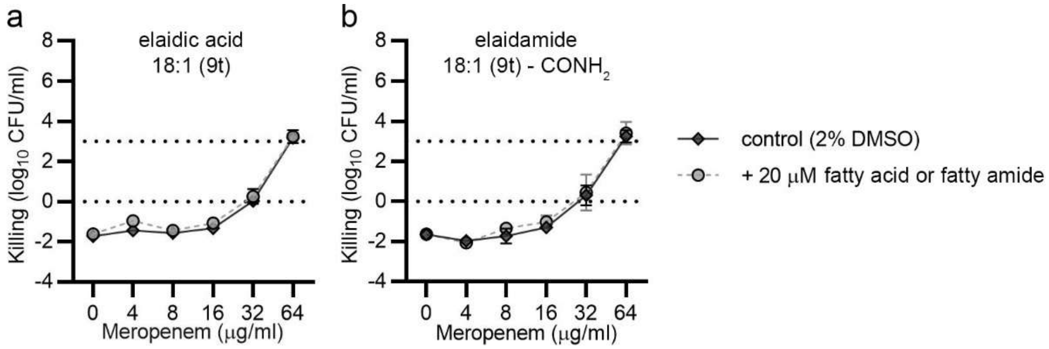


**Fig. S6.** Trans-unsaturated fatty acids and trans-unsaturated fatty amides do not affect the susceptibility of carbapenem-resistant *P. aeruginosa* M6A146 to meropenem. Meropenem susceptibility of *P. aeruginosa* M6A146 in (a) elaidic acid, (b) elaidamide, Points and error bars represent the means and standard deviations of ≥3 independent biological replicates. There were no significant differences when controls and fatty acid- or fatty amide-treated groups were compared using a two-tailed Student's t-test with the Holm-Sidak multiple test correction. Horizontal dotted lines at 0 and 3 $log_{10}$ CFU/mL denote inhibitory and bactericidal thresholds, respectively.

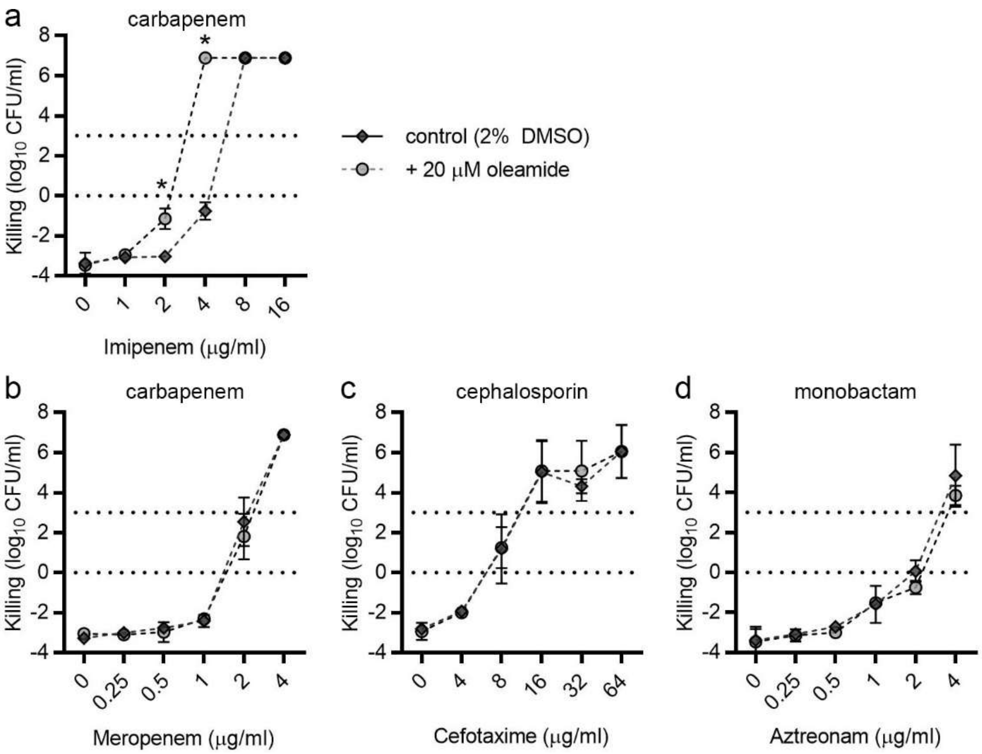


**Fig. S7.** Oleamide potentiates imipenem but not other β-lactam antibiotics against laboratory *P. aeruginosa* PAO1. Susceptibility of *P. aeruginosa* PAO1 to (a) imipenem, (b) meropenem, (c) cefotaxime, and d) aztreonam with and without the addition of 20 µM oleamide. Controls contained 2% DMSO. Points and error bars represent the means and standard deviations of ≥3 independent biological replicates. Significant differences were calculated using a two-tailed Student's t-test with the Holm-Sidak multiple test correction (* denotes $p < 0.05$). Horizontal dotted lines at 0 and 3 $log_{10}$ CFU/mL denote inhibitory and bactericidal thresholds, respectively.

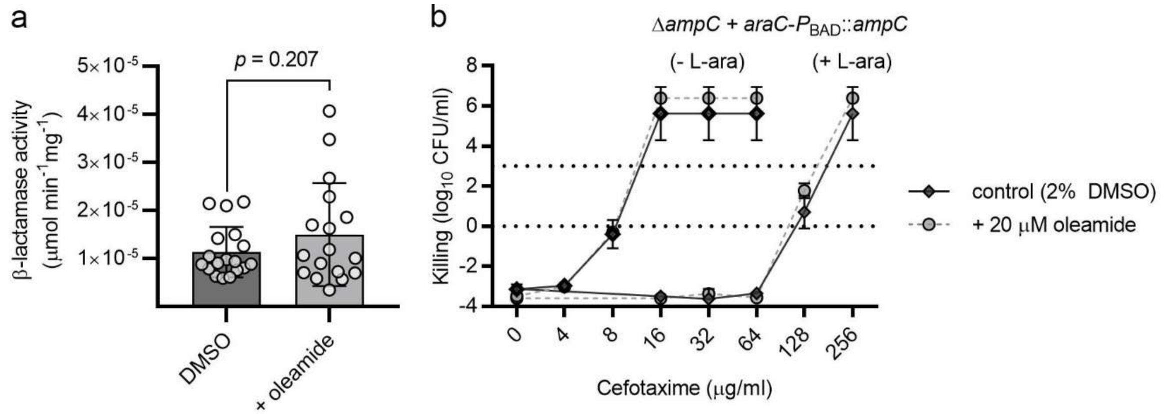


**Fig. S8.** β-lactamases are not the target of oleamide. (a) β-lactamase activity in *P. aeruginosa* M6A146 periplasmic fractions with and without the addition of exogenous oleamide. Circles represent technical replicates from 4 independent biological replicates. There was no significant difference when controls and oleamide treated groups were compared with Student's t-test. (b) Cefotaxime susceptibility of *P. aeruginosa* PAO1 engineered to overexpress AmpC with L-(+)-arabinose (L-ara) induction. Points and error bars represent the means and standard deviations of ≥3 independent biological replicates. There were no significant differences when controls and oleamide treated groups were compared using a two-tailed Student's t-test with the Holm-Sidak multiple test correction. Controls contained 2% DMSO. Horizontal dotted lines at 0 and 3 $\log_{10}$ CFU/mL denote inhibitory and bactericidal thresholds, respectively.

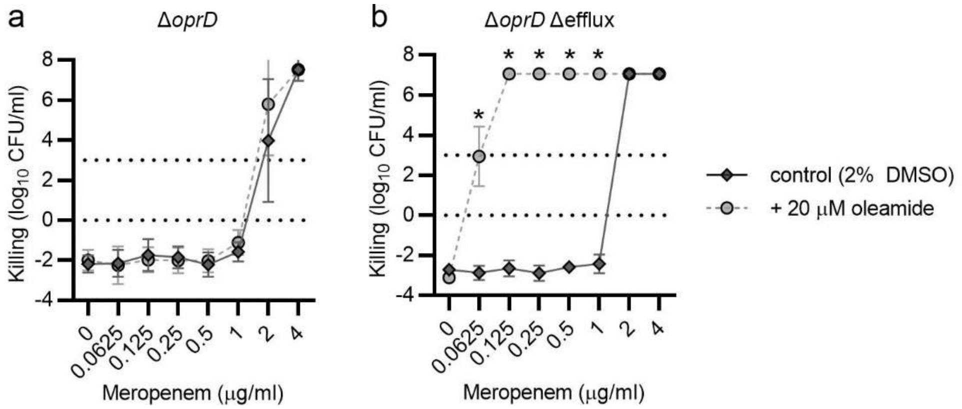


**Fig. S9.** Efflux is a key determinant of meropenem resistance in *P. aeruginosa* PAO1. Susceptibility of *P. aeruginosa* PAO1 (a) Δ*oprD* and (b) Δ*oprD* Δefflux mutants to meropenem with and without the addition of oleamide. Points and error bars represent the means and standard deviations (SDs) of 4 independent biological replicates. Significant differences were calculated using a two-tailed Student's t-test with the Holm-Sidak multiple test correction (* denotes $p < 0.05$). Horizontal dotted lines at 0 and 3 $log_{10}$ CFU/mL denote inhibitory and bactericidal thresholds, respectively.

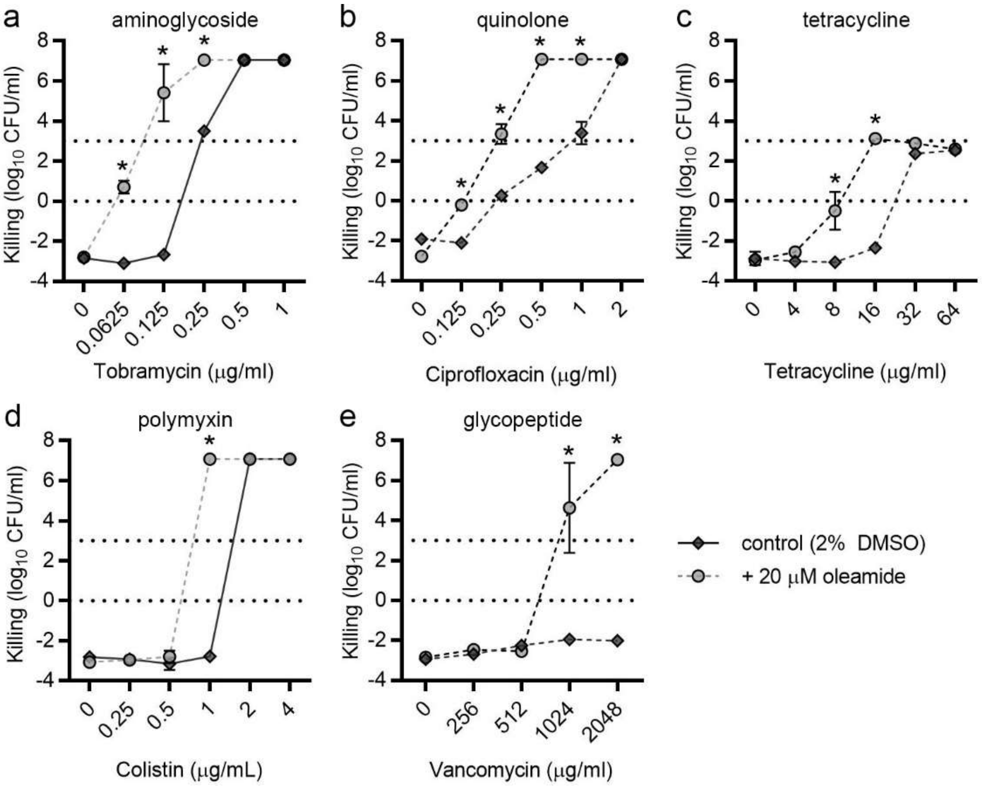


**Fig. S10.** Oleamide potentiates multiple other classes of antibiotics against *P. aeruginosa* PAO1. Susceptibility of *P. aeruginosa* PAO1 to (a) tobramycin, (b) ciprofloxacin, (c) tetracycline, d) colistin, and e) vancomycin, with and without the addition of 20 µM oleamide. Controls contained 2% DMSO. Points and error bars represent the means and standard deviations of ≥3 independent biological replicates. Significant differences were calculated using a two-tailed Student's t-test with the Holm-Sidak multiple test correction (* denotes $p < 0.05$). Horizontal dotted lines at 0 and 3 $log_{10}$ CFU/mL denote inhibitory and bactericidal thresholds, respectively.

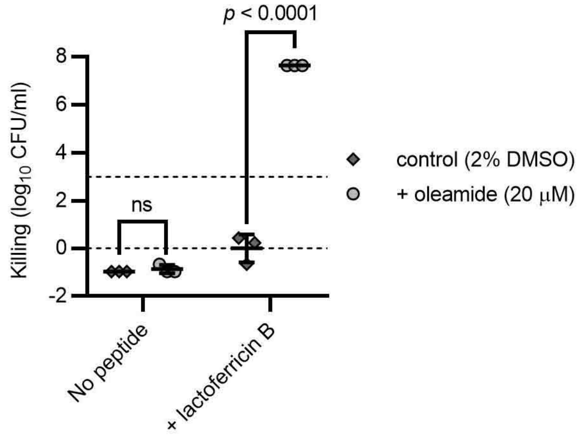


**Fig. S11.** Oleamide significantly augments the bactericidal activity of the lactoferricin antimicrobial peptide. Susceptibility of *P. aeruginosa* PAO1 with and without the addition of 250 µM lactoferricin B. Datum points, lines and error bars represent single biological replicates, means, and standard deviations (SDs), respectively. Significant differences were calculated using a two-tailed Student’s t-test.

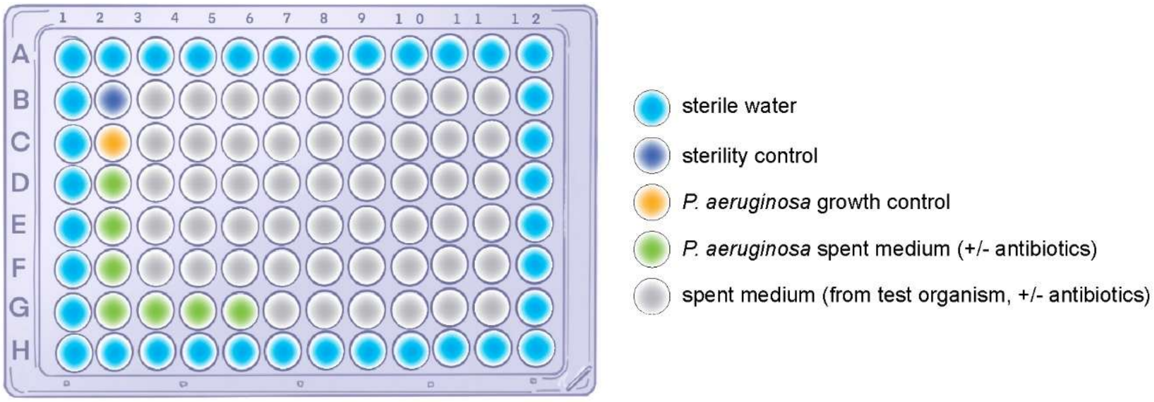


**Fig. S12.** Configuration of content in microplates for medium throughput screening. Microplates configured to assess antibiosis of spent media did not contain antibiotics. By contrast those configured to assess antibiotic potentiation contained antibiotics (2 µg/mL MEM in the context of this study). See Methods for additional details.

**Table S1.** Reference, laboratory, engineered and/or previously isolated bacterial strains used in this study.

| Strain | Genotype, description or relevant characteristics [a] | Source |
|---|---|---|
| *Pseudomonas aeruginosa* | | |
| JJH0 | PAO1 wild type strain from Dr. Colin Mannoil; genome re-sequenced | [2] |
| RP146 | JJH0 Δ*ampC* | This study |
| RP20 | RP146 attTn7::miniTn7T2.1-Gm-GW-*araC*-$P_{BAD}$::*ampC*, GEN$^{r}$ | This study |
| M6A146 | Sputum isolate from a cystic fibrosis patient, MEM$^{r}$ | This study |
| ATCC 27853 | Wild type, reference strain for antibiotic susceptibility testing | ATCC |
| MB5919 | PAO1 wild type strain | [3] |
| MB5919 *lptD4213* | MB5919 harboring *lptD4213* allele, encoding LptD$^{\Delta330\text{-}352}$ | [3] |
| MB5890 | MB5919 Δ(*mexAB-oprM*)::FRT Δ(*mexCD-oprJ*)::FRT Δ*mexXY*::FRT Δ*mexJKL*::FRT Δ(*mexHI-opmD*)::FRT Δ(*opmH*)::FRT | [3,4] |
| MB6476 | MB5890 Δ*oprD* | [3] |
| MB6477 | MB5919 Δ*oprD* | [3] |
| AFC-7 Kenya | Clinical isolate, produces VIM-2 metallo-β-lactamase, MEM$^{r}$ | [5] |
| CF39S | Cystic fibrosis isolate (*tdcA*$^{+}$) | [6] |
| JJH1351 | Bloodstream isolate, produces imipenemase, MEM$^{r}$ | This study |
| JJH1352 | Bloodstream isolate, produces VIM-7 metallo-β-lactamase, MEM$^{r}$ | This study |
| JJH1353 | Bloodstream isolate, produces Guiana extended-spectrum 5 (GES-5) β-lactamase, MEM$^{r}$ | This study |
| JJH1355 | Cystic fibrosis isolate, OprD$^{W65*}$, MEM$^{r}$ | This study |
| JJH1357 | Cystic fibrosis isolate, OprD$^{W277*}$, MEM$^{r}$ | This study |
| JJH1366 | Cystic fibrosis isolate, OprD$^{W138*}$, MEM$^{r}$ | This study |
| *Acenitobacter baumanii* | | |
| HC44 | Clinical isolate | H. Ceri |

| | | |
|---|---|---|
| *Staphylococcus aureus* | | |
| ATCC 13709 | Wild type, reference strain for antibiotic susceptibility testing, $MET^S$ | ATCC |
| *Escherichia coli* | | |
| DH5α | Cloning strain | NEB |
| S17.1 ($\lambda_{pir}$) | Mobilization host for conjugation | |
| ATCC 25922 | Wild type, reference strain for antibiotic susceptibility testing | ATCC |

[a]Abbreviations for antibiotic selection: GEN, Gentamycin; MEM, meropenem; MET, methicillin.

**Table S2.** Plasmids used in this study.

| Plasmid | Genotype, description, or relevant characteristics [a] | Source |
|---|---|---|
| **Cloning, template, and helper plasmids** | | |
| pJN105 | Cloning vector with *araC*-$P_{BAD}$ expression cassette, $GEN^r$ | [7] |
| pTNS3 | Helper plasmid for miniTn7 mutagenesis, $AMP^r$ | [8] |
| **Allelic exchange vectors** | | |
| pDONRPEX18Gm | Allelic exchange vector with Gateway donor site, *sacB*, $GEN^r$, $CHL^r$ | [2,9] |
| pRP23 | pENTERPEX18Gm::Δ*ampC*, $GEN^r$ | This study |
| **miniTn7 vectors for site-specific chromosomal insertions** | | |
| pUC18-miniTn7T2.1-Gm-GW | miniTn7 with transcriptional terminators at both ends of the transposon; Gateway destination vector $CHL^r$, $GEN^r$, $Ap^r$ | [10] |
| pRP1 | pUC18-miniTn7T2.1-Gm-GW--*araC*-$P_{BAD}$::*ampC*, $Gm^r$, $Ap^r$ | This study |

[a]Abbreviations for antibiotic selection: AMP, ampicillin; CHL, chloramphenicol, GEN, Gentamycin.

**Table S3.** Primers.

| Oligonucleotide | DNA sequence[a] |
|---|---|
| **DNA sequencing** | |
| 27F | AGA GTT TGA TCC TGG CTC AG |
| 926R | CCG TCA ATT CMT TTR AGT |
| 1492R | TAC CTT GTT ACG ACT T |
| ITS1-F | CTT GGT CAT TTA GAG GAA GTA A |
| ITS4 | TCC TCC GCT TAT TGA TAT GC |
| oJJH367_M13-Universal-F | GTA AAA CGA CGG CCA G |
| oJJH368_M13-Universal-R | CAG GAA ACA GCT ATG AC |
| **Construction of allelic exchange vectors** | |
| oRP_43 | GGG G**AC AAG TTT GTA CAA AAA AGC AGG CTA C**CC GTG AAG GTT CCG ACC G |
| oRP_44 | *CGC TGA GGA TGG CGT AGG CGA TCT TC*G CAG GGG AAT CTG GTA TCG C |
| oRP_45 | GAA GAT CGC CTA CGC CAT CC |
| oRP_46 | GGG G**AC CAC TTT GTA CAA GAA AGC TGG GTA** GTC TTG AGC GAC ATC CTC TTC |
| **Construction of miniTn7 vectors** | |
| oRP_51_ampC-F1 | GCG AAT TCC TGC AGC CCG GGG GAT CCA CTG AGG AGG ATA TTC ATG CGC GAT ACC AGA TTC |
| oRP_52_ampC-R1 | TGG TTG GCC TGC AAG GCC TTC GCG AGG TAC CTT CAG CGC TTC AGC GGC AC |
| oHA229_miniTn7_fwr_pBAD | GAT CCC CCG GGC TGC AGG AAT TCC TCG AGA TTA TGA CAA CTT GAC GGC TAC AT |
| oHA230_miniTn7_rev_pBAD | AGT GGA TCC CCC GGG CTG CAG GAA TTC GCT |
| oRP_92_pUC18T-miniTn7-Gm-GW_fwd | GTA CCT CGC GAA GGC C |
| oRP_93_pUC18T-miniTn7-Gm-GW_rev | TCT CGA GGA ATT CCT GCA G |

[a]*Italics* denote a region of complementarity for SOE-PCR; underlined sequence denotes a region specific to the targeted DNA amplicon; **bold** denotes an attachment (*att*) or restriction site for cloning.

**Table S4.** Culture media compositions.

| Culture medium (abbreviation) | Components (per liter of ultrapure water) | Description and/or material source(s) |
|---|---|---|
| 1/10 869[11] | 0.035 g calcium chloride, 0.5 g sodium chloride, 1.0 g tryptone, 0.5 g yeast extract, 15 g agar, 0.1 g/L glucose (after autoclaving) | An undefined medium that is used primarily to isolate plant endophytes.<br>Calcium chloride, Fisher, Ref C-79 (500g) Lot: 720738<br>Sodium chloride, VWR, Ref BDH9286-2.5KG Lot: 20B2556917<br>Tryptone, Bacto Ref J859-500G Lot: 22A24566376<br>Yeast extract, Bacto Ref 212750-500G Lot: 4272947<br>Granulated agar, Difco BD-500G, Ref 214530<br>Glucose, Fisher Bioreagents, Ref BP350-1, Lot: 170461 |
| Bile esculin agar (BEA)[12] | 5.0 g gelatin peptone, 3.0 g beef extract, 40 g bile, 1.0 g esculin, 0.5 g ferric citrate, 15 g agar | For cultivation of group D streptococci and enterococci but inhibits Gram-negative bacteria<br>Gelatin peptone, Criterion, Ref C6541 Lot: 14238<br>Meat extract, Microbiology, Lot: VM659479 431<br>Bile, Sigma, Ref B-8381, Lot: 110H0215<br>Difco labs, Ref 0158-12-6 Lot: 83359JA<br>Ferric citrate, Sigma, Ref F3388 (250g) Lot: SLBL6061V<br>Granulated agar, Difco BD-500G, Ref 214530 |
| Brain heart infusion (BHI) | 200 g [7.7 g/L calf brain infusion], 250 g [9.8 g/L beef heart infusion], 10 g proteose peptone, 2.0 g dextrose, 5.0 g sodium chloride, 2.5 g disodium phosphate, 15 g agar | A general-purpose medium used for aerobic cultivation of many fastidious bacteria, fungi and yeast, including pathogens<br>BBL BD-500G, Ref 211059 Lot: 1159375. |
| Brain heart infusion III + Supplement Set A (BHI-A)[13] | BHI + Supplement A (Table S2) | To capture diverse intestinal microbiota |
| Brain heart infusion II + Supplement Set B (BHI-B)[13] | BHI + Supplement B (Table S2) | To capture diverse intestinal microbiota |
| Brain heart infusion IV + Supplement Set A & B (BHI-AB)[13] | BHI + Supplement A + Supplement B (Table S2) | To capture diverse intestinal microbiota |

**Table S4.** Culture media compositions (continued).

| Culture medium | Components (per liter of ultrapure water) | Description and/or material source |
|---|---|---|
| Brewer's anaerobic agar (BAA)[13] | 5.0 g pancreatic digest of casein, 10.0 g proteose peptone no. 3, 5.0 g yeast extract, 10 g dextrose, 5 g sodium chloride, 20 g agar, 2.0 g sodium thioglycolate, 1.0 g sodium formaldehyde sulfoxylate, 2.0 mg resazurin | This non-selective medium is used to isolate facultative and obligate anaerobes.<br>Difco BG 500G, Ref 227920 Lot: 4310891 |
| Brucella blood agar + 5% sheep's blood | 10 g pancreatic digest of casein, 10 g peptic digest of animal tissue, 1.0 g dextrose, 2.0 g yeast extract, 5.0 g sodium chloride, 0.1 g sodium bisulfite and 15 g agar<br>+ 5% sheep's blood (Table S2) | Selective medium for isolation of fastidious, obligately anaerobic organisms<br>Himedia 500G, Ref M823 Lot: 0000188103 |
| Chocolate blood agar | 7.5 g tryptone, 7.5 g HM extract, 5.0 g sodium chloride, 4.0 g dipotassium hydrogen phosphate, 1.0 g corn starch, 1.0 g potassium dihydrogen phosphate, 12.0 g agar | A nonselective, enriched medium is used for fastidious organisms like *Neisseria* sp., *Haemophilus* sp. and *Gardnerella vaginalis*.<br>Himedia 500G, Ref M1548 Lot: 0000301775 |
| Columbia blood agar + 5% sheep's blood (CBA) | 10 g pancreatic digest of casein, 3.5 g proteose peptone, 5.0 g yeast extract, 500 g 3.0 g/L beef heart infusion, 1.0 g corn starch, 5.0 g sodium chloride, 15 g agar | A multi-purpose medium used to grow fastidious and non-fastidious bacteria. Does not support the growth of *Haemophilus*<br>Difco BD-500G, Ref 279240 Lot: 7297711 |
| Cooked meat medium from dehydrated meat extracts (Meat) | 98 g beef heart, 20 g proteose peptone, 2.0 g dextrose, 5.0 g sodium chloride | An enriched medium for isolation of anaerobic bacteria such as *Clostridium sp*.<br>Himedia, Ref M1040-500G Lot: 0000348605 |
| Cooke rose bengal agar (CRB) | 5.0 g papaic digest of soyabean meal, 10 g dextrose, 1.0 g monopotassium phosphate, 0.5 g magnesium sulphate, 0.035 g rose bengal and 20 g agar | This selective medium permits growth of fungi.<br>DifcoTM, BD-500G, Ref 270310 Lot: 4307881 |
| Desoxycholate (DOC) | 10 g peptone, 10 g lactose, 1g sodium desoxycholate, 5g sodium chloride, 2.0 g dipotassium phosphate, 1g ferric ammonium citrate, 1.0 g sodium citrate, 15 g agar and 0.03 g neutral red | A selective and differential medium to grow of Gram-negative, enteric bacteria.<br>Difco BD-500G, Ref 227310 Lot: 5019893 |
| Fastidious anaerobic agar (FAA) | 23 g peptone, 5.0 g sodium chloride, 1.0 g soluble starch, 0.4 g sodium bicarbonate, 1.0 g glucose, 1.0 g sodium pyruvate, 0.5 g L-cysteine HCl•$H_20$, 0.25 g sodium pyrophosphate, 1.0 g L-arginine, 0.5 g sodium succinate, 0.01 g hemin, 0.001 g vitamin K and 12 g agar. | An isolation medium for anaerobic bacteria, including fastidious organisms.<br>Acumedia 7531B-2kg, Ref 02-37286-00C Lot: 109111C |

**Table S4.** Culture media compositions (continued).

| Culture medium | Components (per liter of ultrapure water) | Description and/or material source |
|---|---|---|
| Kligler iron agar (KIA) | 10 g pancreatic digest of casein, 10 g peptic digest of animal tissue, 10 g lactose, 1.0 g dextrose, 5.0 g sodium chloride, 0.5 g ferric ammonium citrate, 0.5 g sodium thiosulfate, 15 g agar, 25 mg phenol red | This medium is used to differentiate bacteria on the basis of hydrogen sulfide production.<br>BBL BD-500G, Ref 211317 Lot: 4124934 |
| Lysogeny broth agar (LB) | 10 g tryptone, 5.0 g yeast extract, 5.0 g sodium chloride and 15 g granulated agar | A general-purpose medium for growth and cultivation of bacteria, particularly members of Enterobacteriacea.<br>Tryptone, Bacto Ref J859-500G Lot: 22A24566376<br>Yeast extract, Bacto Ref 212750-500G Lot: 4272947<br>Sodium chloride, VWR Chemicals Ref BDH9286-2.5KG Lot: 20B2556917<br>Granulated agar, Difco BD-500G, Ref 214530 |
| M9 minimal medium (M9) | 200 mL of 5 × M9 salt solution [33.9 g anhydrous disodium phosphate, 15 g monopotassium phosphate, 2.5 g sodium chloride, 5 g ammonium chloride], 0.4% glucose, 2 mM $MgSO_4$, 0.1 mM $CaCl_2$ and 15 g agar | A selection medium for plasmid-containing strains of *E. coli* with the ability to produce essential amino acids or vitamins.<br>Difco BD-500G, Ref 248510 Lot: 8023596 |
| M9 minimal media + insulin[3]<br>(M9 + insulin) | M9 + 1 g/L insulin | To capture diverse intestinal microbiota.<br>Sigma-Aldrich, 1-6634, CAS 11070-73-8 Lot: 40H00235 |
| M9 minimal media + mucin[13]<br>(M9 + mucin) | M9 + 0.5 g/L mucin | To capture diverse intestinal microbiota<br>Sigma-Aldrich, M1778-10g, Lot: 68H7480 |
| M9 minimal media + starch[13]<br>(M9 + starch) | M9 + 0.5 g/L starch | To capture diverse intestinal microbiota<br>Fisher Scientific, S-516-1Lb, Ref 79013 Lot: 756146 |
| M17 minimal medium (M17) | 5.0 g pancreatic digest of casein, 5.0 g soy peptone, 5.0 g beef extract, 2.5 g yeast extract, 0.5 g ascorbic acid, 0.25 g magnesium sulfate, 19 g disodium-β-glycerophosphate, 11 g agar | This growth medium is used to enumerate lactic streptococci from dairy products.<br>Difco BD-500G, Ref 218561 Lot: 5054747 |
| MacConkey agar (MCA) | 17 g pancreatic digest of gelatin, 3.0 g peptones (meat and casein), 10 g lactose, 1.5 g bile salts no. 3, 5.0 g sodium chloride, 13.5 g agar, 0.03 g neutral red, 1 mg crystal violet | This medium is selective for growth of Gram-negative enteric bacteria and differentiates based on their ability to ferment lactose.<br>Difco BD-500G, Ref 211387 Lot: 6313690 |

**Table S4.** Culture media compositions (continued).

| Culture medium | Components (per liter of ultrapure water) | Description and/or material source |
|---|---|---|
| Marine broth agar (MBA) | 5.0 g peptone, 1.0 g yeast extract, 0.1 g ferric citrate, 19.45 g sodium chloride, 5.9 g magnesium chloride, 3.24 g magnesium sulfate, 1.8 g calcium chloride, 0.55 g potassium chloride, 0.16 g sodium bicarbonate, 0.08 g potassium bromide, 0.034 g strontium chloride, 0.022 g boric acid, 0.004 g sodium silicate, 0.0024 g sodium fluoride, 0.0016 g ammonium nitrate, 0.008 g disodium phosphate and 15 g agar | To grow heterotrophic bacteria from aquatic environments like *Vibrio sp*.<br>Difco BD-500G, Ref 212185 Lot: 9070997 |
| No salt lysogeny broth agar (NSLB)[4] | 10 g tryptone, 5.0 g yeast extract, 15 g agar | A general, nutrient-rich medium for bacterial culturing. Most commonly supports *E. coli* growth.<br>Tryptone, Bacto Ref J859-500G Lot: 22A24566376<br>Yeast extract, Bacto Ref 212750-500G Lot: 4272947<br>Granulated agar, Difco BD-500G, Ref 214530 |
| No salt lysogeny broth agar + 5-15% sucrose (NSLB + 5-15% sucrose) | 10 g tryptone, 5.0 g yeast extract, 5.0 to 15 g sucrose (as required) and 15 g agar | Tryptone, Bacto Ref J859-500G Lot: 22A24566376<br>Yeast extract, Bacto Ref 212750-500G Lot: 4272947<br>Sucrose, Fisher Bioreagents, BP220-212 Lot: 208132<br>Granulated agar, Difco BD-500G, Ref 214530 |
| Nutrient agar | 3.0 g beef extract, 5.0 g peptone and 15 g agar | A general-purpose medium used to grow many non-fastidious microorganisms.<br>Difco BD-500G, Ref 213000 Lot: 5027887 |
| Phenylethyl alcohol agar + 5% sheep's blood (PEA)[13] | 15 g pancreatic digest of casein, 5.0 g papaic digest of soybean meal, 5.0 g sodium chloride, 2.5 g β-phenylethyl alcohol, 15 g agar | A selective medium supporting the growth of gram-positive cocci but inhibits gram-negative and fungi.<br>BBL, BD-500G, Ref 211539 Lot: 4303910 |
| Plate count medium (PC) | 5.0 g pancreatic digest of casein, 2.5 g yeast extract, 1.0 g dextrose and 15 g agar | This nonselective medium is used to count bacterial colonies grown on agar plates.<br>Difco BD-500G, Ref 275120 Lot: 4139667 |
| Pseudomonas isolation medium (PIA) | 20 g peptone, 1.4 g magnesium chloride, 10 g potassium sulfate, 25 mg Irgasan, 13.6 g agar | A selective differential medium for cultivation and isolation of *Pseudomonas sp*.<br>Difco BD-500G, Ref 292710 Lot: 7009633 |

**Table S4.** Culture media compositions (continued).

| Culture medium | Components (per liter of ultrapure water) | Description and/or material source |
|---|---|---|
| Reasoner's 2A agar (R2A) | 0.5 g yeast extract, 0.5 g proteose peptone no. 3, 0.5 g casamino acids, 0.5 g dextrose, 0.5 g soluble starch, 0.3 g sodium pyruvate, 0.3 g dipotassium phosphate, 0.05 g magnesium sulfate and 15 g agar | Low nutrient medium that enables growth of stressed and chlorine-tolerant bacteria from potable water after long incubation periods at lower temperatures<br>Difco BD-500G, Ref 218263 Lot: 7151674 |
| Reinforced Clostridial agar (RCA) | 5.0 g tryptone, 5.0 g peptone, 8.0 g beef extract, 1.0 g yeast extract, 0.5 g L-cysteine HCl, 1.0 g starch, 1.0 g dextrose, 5.0 g sodium acetate, 0.5 g sodium bisulfite, 0.5 g ferric ammonium citrate, 2 mg resazurin, 15 g agar | To grow sulfite-reducing clostridia<br>EMD TM 500G, Lot: VM036511 |
| Staphylococcus isolation medium (SIA) | 10 g pancreatic digest of casein, 2.5 g yeast extract, 2.0 g lactose, 10 g mannitol, 5.0 g dipotassium phosphate, 75 g sodium chloride | A selective medium used to cultivate enterotoxigenic *Staphylococcus* species.<br>BD Difco BD-500G, Ref 264920 Lot: 6277940 |
| Supplemented BHI (sBHI 2.0)[14] | BHI + Supplement 2.0 (Table S2) | Anaerobic bacterial growth medium for isolation of vaginal microbes. |
| Tryptic soy agar + Supplement A & B + yeast (TSY)[13] | 200 g [7.7 g/L calf brain infusion], 250 g [9.8 g/L beef heart infusion], 10 g proteose peptone, 2.0 g dextrose, 5.0 g sodium chloride, 2.5 g disodium phosphate, and 15 g agar + Supplement A + Supplement B + 5.0 g yeast extract | To capture diverse intestinal microbiota<br>Difco BD-500g, Ref 236950 Lot: 2158089<br>Yeast extract, Bacto Ref 212750-500G Lot: 4272947 |

**Table S5.** Supplements.

| Supplement | Components (per liter of ultrapure water) | Description and/or material source |
|---|---|---|
| 5% sheep's blood | --- | Life Sciences Animal Research Center, University of Calgary |
| 5% fetal bovine serum | --- | Life Sciences Animal Research Center, University of Calgary |
| Supplement A[13] | 0.5 g/L L-cysteine hydrocholate hydrate<br>10 mg/L hemin<br>1 mg/L vitamin K | Sigma-Aldrich, C-7880-100G, Ref 7048-04-6 Lot: 090K06041<br>Sigma-Aldrich, H9039-1G, CAS 16009-13-5 Lot: SLBP5720V<br>Sigma-Aldrich, 95271-1G, CAS 84-80-0 |
| Supplement B[13] | 10 mg/L colistin sulphate<br>5 mg/L nalidixic acid | Sigma-Aldrich, C4461-1G, STB2006006 Batch 0000118405<br>Sigma-Aldrich, N8878-5G, CAS 389-08-2 Lot: BCBR7632V |
| Supplement 2.0[14] | 8.0 g/L glucose,<br>10 g/L yeast extract,<br>1.0 g/L starch<br>20 g/L gelatin | Fisher Bioreagents, Ref BP350-1 (1kg) Lot: 170461<br>Bacto Ref 212750-500G Lot: 4272947<br>Fisher Scientific Company, S-516, Ref 79013 Lot: 756146<br>Sigma, Ref G-6269 (100g) Lot: 97F0748 |
